# Single particle dynamics of protein aggregation and disaggregation in the presence of the sHsp proteins IbpAB

**DOI:** 10.1101/2025.05.07.652671

**Authors:** Andrew Roth, Jason Puchalla, Hays S. Rye

**Author notes:** Correspondence should be addressed to H. S. R.

## Abstract

The small heat shock proteins (sHsps) are a key class of molecular chaperones that can inhibit protein aggregation and enhance protein recovery from aggregates. However, the mechanisms sHsps employ to carry out these roles are not well understood, in part because the highly heterogeneous and dynamic particles they form with aggregating proteins are difficult to study with traditional biophysical tools. Here we have applied a novel single particle fluorescence technique known as Burst Analysis Spectroscopy (BAS) to the study of the *E. coli* sHsps IbpA and IbpB (IbpAB). We show that in the presence of IbpAB, two different model proteins converge toward similar, limited aggregate particle size distributions. Additionally, while IbpAB dramatically accelerates the disassembly of protein aggregates by the bacterial KJEB bi-chaperone disaggregase, this enhancement does not appear to be strongly influenced by aggregate particle size. Rather, it is the ability of IbpAB to alter aggregate structure during particle formation that appears to be essential for stimulated disassembly. These observations support a model of aggregate recognition by IbpAB that is not only highly adaptable but capable of shaping aggregate particles into a specialized range of physical properties that are necessary for efficient protein disaggregation.

## INTRODUCTION

Protein folding and assembly can be error prone and inefficient, particularly in the crowded interior of a living cell (1–3). To ensure that folding succeeds more often than it fails, several classes of facilitator proteins, known as molecular chaperones, arose early in cellular evolution (4, 5). In addition to supporting basic protein homeostasis, molecular chaperones also play key roles in mitigating the impacts of environmental stress (6, 7). A variety of perturbations, including elevated temperature, altered pH, and oxidation damage can cause protein misfolding, aggregation and cellular dysfunction. Aberrant protein folding and aggregation have been implicated in aging, as well as a wide variety of diseases, including cataracts, cardiomyopathy, and neurodegenerative disorders (8, 9).

One of the oldest families of molecular chaperones, and an essential first line of defense in the cellular response to protein stress, are the small heat shock proteins (sHsp) (4, 10–12). Among a number of important functions, the sHsps have a primary role in addressing protein aggregation, acting to both inhibit aggregation and facilitate disaggregation and recovery (13–15). Unlike many other molecular chaperone families, the sHsps possess no ATP binding sites and their interactions with client proteins are thus ATP-independent (16). While the sHsps have diversified in both size and function throughout evolutionary history, they share several common features, including a conserved, central α-crystallin domain (ACD) and flexible, mostly unstructured, N- and C-terminal domains (15, 17). Another signature feature of the sHsps is their ability to form large and dynamic oligomeric complexes, which can vary widely in configuration between the different sHsp sub-families (18–20). Both the ACD and N- and C-terminal domains play important roles in the formation of both small oligomers and larger complexes, with the detailed nature of the interactions varying among sHsp types (13, 14). A common feature of sHsps is their tendency to form large, inactive oligomers under non-stress conditions that disassemble into active subunits, often dimers, following an environmental stress like heat shock (21–23).

While many aspects of sHsp structure and function have been well characterized, how they interact with aggregating proteins and facilitate aggregate disassembly remains poorly understood. In general, once activated, sHsps form heterogeneous co-complexes with non-native, aggregation-prone proteins, which can span a range of particle sizes, stoichiometries and physical properties (10, 12, 24). In many cases, sHsps reduce the average size of aggregate particles that form in their presence, relative to those that form in their absence (10, 12, 25, 26). Because uninhibited aggregate growth is associated with lower solubility, increase average particle size and lower active protein recovery, it is generally thought that sHsps act to block formation of large, difficult to process aggregate particles (13–15). Several models have been proposed to explain how sHsps accomplish this task. In one model, sHsps form dynamic, shell-like structures around the exterior of aggregate particles, physically blocking ongoing aggregate growth (27). In another model, sHsps function primarily to alter the packing of non-native protein subunits in an aggregate, using features of the ACD and N-terminal domains to directly incorporate into the aggregate particle structure (28, 29). It has also been proposed that sHsps have evolved to recognize and bind near-native conformations of folding intermediates, reducing their aggregation propensity and improving subsequent reactivation (30–32).

Here, we focus on the *E. coli* sHsps IbpA and IbpB (IbpAB), which form a functional heterodimer (33–35). IbpAB plays an important role in limiting protein aggregation and facilitating aggregate disassembly, in partnership with the bi-chaperone disaggregase of DnaK, DnaJ, GrpE and ClpB (KJEB) (36–39). Despite much progress, most prior studies have employed signatures of native state recovery to measure disaggregation in the presence of IbpAB (e.g, enzymatic reactivation). For many such proteins, disaggregation and refolding are coupled events, with the KJE chaperone system directly involved in both disaggregation and folding (27, 34, 36, 37) This overlap generates ambiguity in the assignment of IbpAB action to specific experimental events. Mechanistic analysis is made even more difficult by the complex and dynamic nature of the large assemblies formed between protein aggregates and IbpAB. Here, we overcome these limitations by applying a single particle fluorescence-based technique called Burst Analysis Spectroscopy (BAS) (40, 41). BAS provides a highly flexible, real-time approach to quantifying the population-resolved kinetics of biological nanoparticle formation and disassembly under minimally perturbed, free solution conditions.

Using BAS, we previously demonstrated that the CO_2_-fixing enzyme RuBisCO from *Rhodospirillum* rubrum is a versatile model for studying protein aggregation, disaggregation and chaperone-mediated folding (40–44). Fully functional fluorescent variants of the enzyme are straightforward to prepare and slight alterations in solution conditions shift the aggregation behavior of RuBisCO between two different pathways, one distinguished by the exclusive population of slow growing (S-type) aggregate particles, the other enriched in particles that display much faster growth, distinctive physical properties and different population distribution changes (F-type) (44). We have also characterized fluorescent versions of the chaperone-dependent prolidase enzyme of *E. coli*., PepQ (45, 46), which displays aggregation behavior that is distinct from that of both S- and F-type RuBisCO aggregates. Here, we utilize BAS, electron microscopy, and fluorescence resonance energy transfer (FRET) to examine how IbpAB impacts the aggregation and disaggregation of RuBisCO and PepQ. We show that IbpAB appears capable of sculpting both S-type and F-type RuBisCO, as well as PepQ aggregates, into particles of a surprisingly similar overall size. Strikingly, the ability of IbpAB to alter the internal structure of protein aggregate particles, rather than the overall size of the particles, appears to be the primary determinant of whether these sHsp proteins can effectively facilitate disaggregation. We further demonstrate that release of IbpAB from aggregate particles appears to be coincident with aggregate particle disassembly itself, an observation that is inconsistent with simple coating models of IbpAB action.

## RESULTS

### IbpAB limits different aggregating proteins to a similar particle size

We first determined whether RuBisCO is a substrate for IbpAB. When heat activated IbpAB is added to aggregating RuBisCO following the S-type pathway (Figure 1A) (44), aggregate particles appear smaller and more numerous by negative stain electron microscopy (Figure 1B). Using an Alexa647-labeled RuBisCO variant, BAS measurements show that IbpAB induces a shift in the aggregate burst distribution to a significantly smaller mean value (Figure 1C-D). Calibration of the intrinsic brightness of the Alexa647-labeled RuBisCO (Figure S1) shows that the most abundant aggregate particles contain 20-40 RuBisCO monomers (20-40 mer), a reduction in particle size of approximately 10-fold, relative to the S-type aggregates alone.

**Figure 1.**
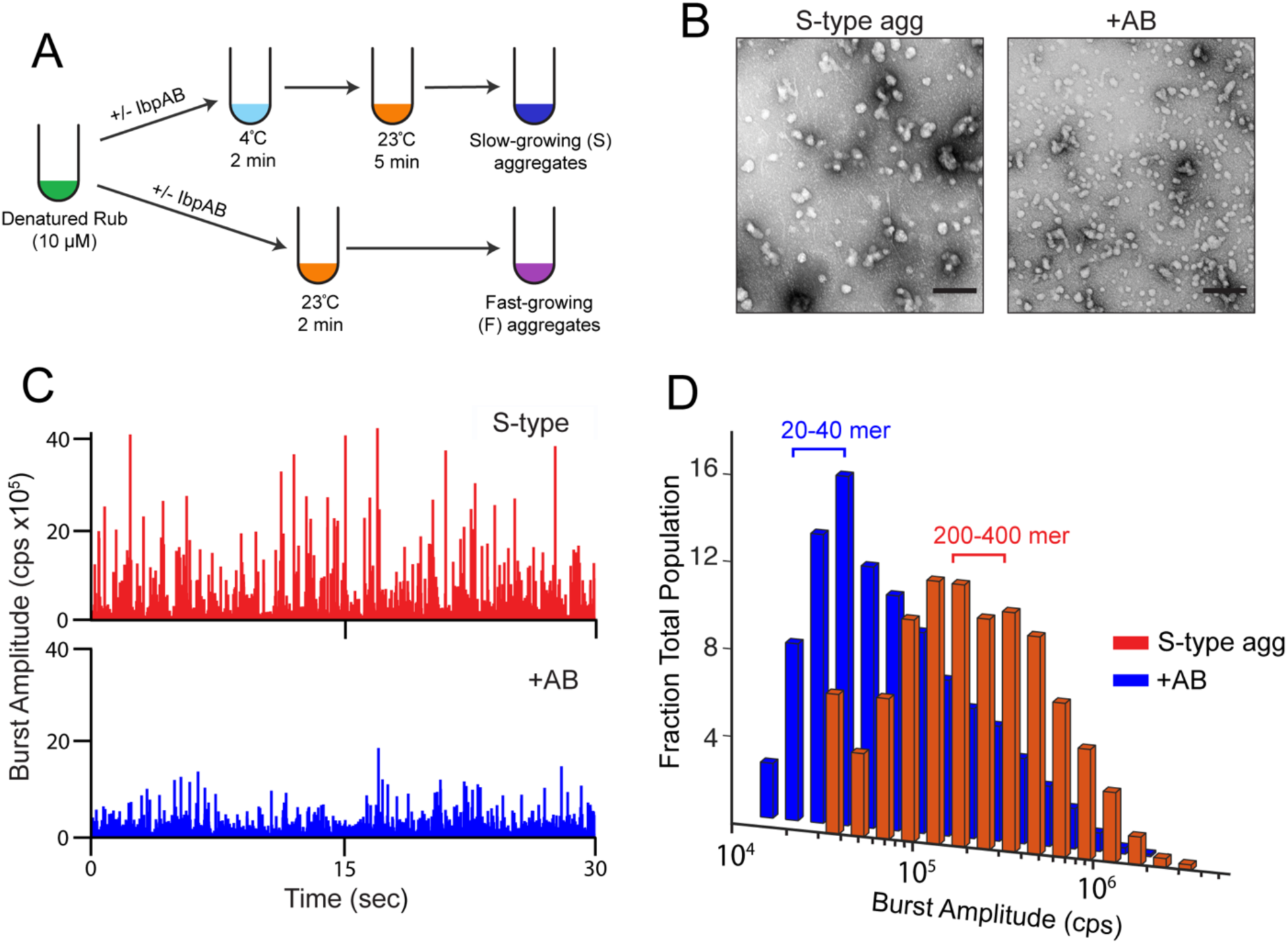
The impact of IbpAB on a protein aggregate distribution can be measured with Burst Analysis Spectroscopy (BAS). (A) Alexa647-labeled RuBisCO (RuBisCO-A647) was denatured in acid urea, then rapidly diluted 50-fold to 200 nM (final monomer concentration) in HKM buffer at either 4 °C or 23 °C, in the presence or absence of heat-activated IbpAB. Samples diluted into cold buffer were then warmed to 23 °C to form S-type aggregates. Direct dilution into a warm (23 °C) buffer results in formation of F-type aggregates. Following a brief incubation period, aggregate growth was halted by dilution to a final monomer concentration of 10 nM. (B) Negative stain electron microscopy images of S-type RuBisCO aggregate particles formed in the presence (1:1 IbpAB:RuBisCO monomer) and absence of heat activated IbpAB (scale bar = 200 nm). Similar behavior was observed with F-type aggregates (not shown). (C) Raw photon history showing fluorescence bursts of S-type RuBisCO-A647 aggregates formed in the presence (*blue*) and absence (*red*) of heat activated wild type IbpAB (50 nM final dimer concentration). (D) Distribution of aggregate particle sizes measured by BAS. The approximate peak of each distribution, shown as the number of RuBisCO monomers per particle, is derived from the measured effective brightness of single Alexa647-labeled RuBisCO monomers incorporated into an aggregate particle (Figure S1 and ref (41)). Each BAS plot is a combination of n = 3, independent experimental replicates.

We next asked whether the large observed reduction in S-type particle size is specific to RuBisCO and the S-type aggregation pathway, or if it is potentially a more general property of IbpAB. To address this question, we examined how IbpAB affects the second, F-type RuBisCO aggregation pathway (Figure 1A) (44), as well as a different substrate protein, the *E. coli* prolidase enzyme PepQ (46). Importantly, we previously characterized a fully functional tetramethyl rhodamine derivative of PepQ (PepQ-TMR) that is ideal for BAS aggregation experiments (45). We employed an approach similar to that described above for RuBisCO to establish the intrinsic brightness of the TMR-labled PepQ monomer (Figure S2).

We used BAS to examine aggregate particle distributions across a range of IbpAB to substrate protein ratios (Figure 2). To more fully characterize the smallest particles in population, we employed higher dilutions (2 to 10-fold) of the aggregating samples than typically used for BAS. This modification permits better characterization of very small amplitude events in the population but requires much longer data collection times. Addition of IbpAB to S-type RuBisCO aggregates results in a concentration-dependent shift in aggregate particle size to smaller values, with the maximal impact observed once the IbpAB dimer to RuBisCO monomer level reaches approximately 5:1 (Figure 2A). At this point, the most abundant aggregate particles are composed of 20-40 RuBisCO monomers. Higher concentrations of IbpAB (up to 10:1) had only a modest additional impact on the observed aggregate distribution, with a small reduction in the larger particle populations and an associated increase in the dominant 20-40 mer sub-population (data not shown). Addition of IbpAB to F-type RuBisCO or PepQ aggregates also results in a concentration-dependent reduction in aggregate particle size (Figure 2B-C). However, despite their distinct physical properties and growth behaviors, both F-type RuBisCO and PepQ aggregates approach a particle size (10-30 mer) that is like that observed with the S-type RuBisCO aggregates. Interestingly, the amount of IbpAB required to populate this smaller particle distribution differs for each aggregate type, with the S-type RuBisCO aggregates requiring the highest level of IbpAB and PepQ requiring the least (Figure 2).

**Figure 2.**
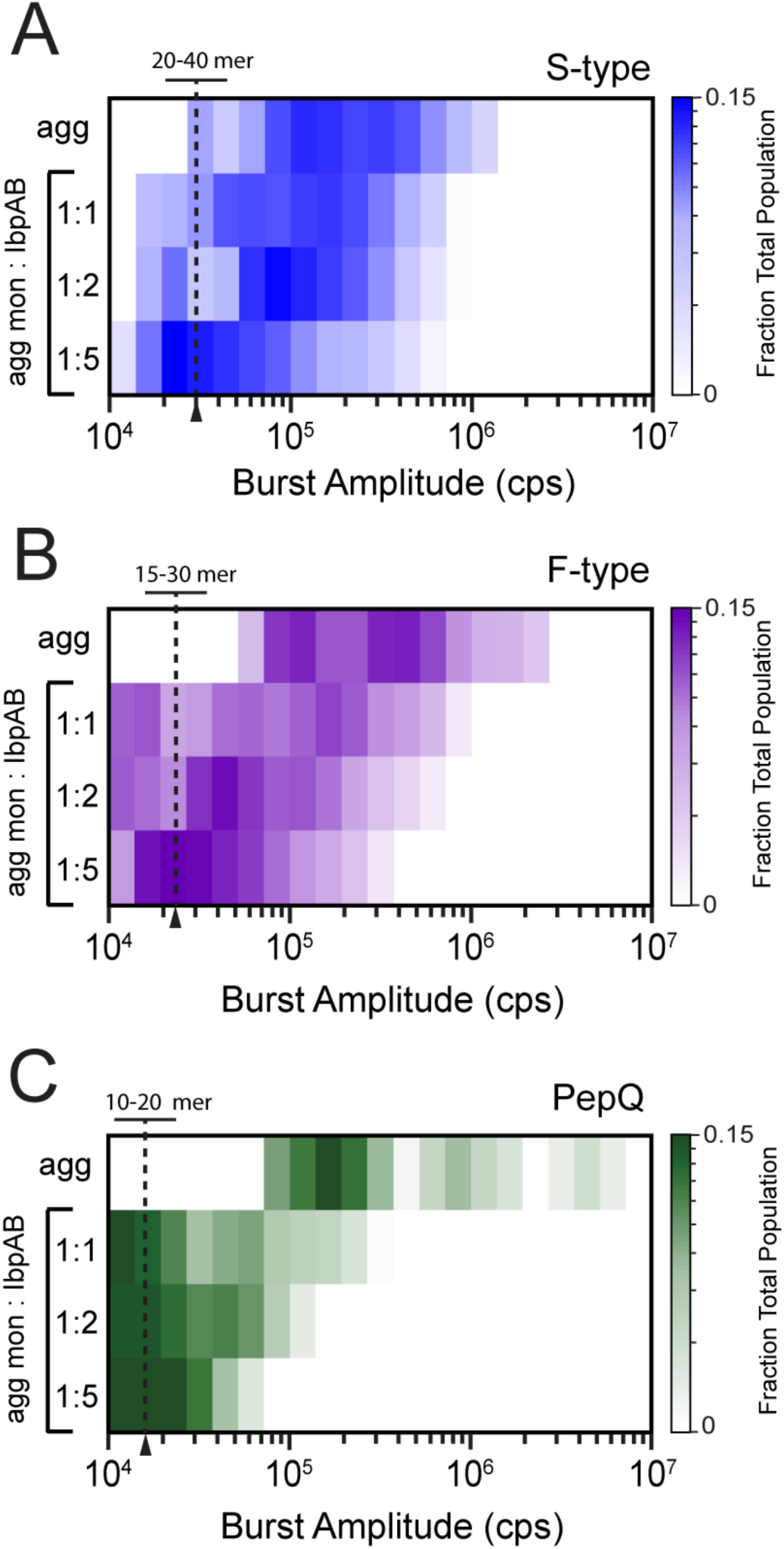
IbpAB restricts the aggregation of different proteins to a similar and limited particle size range. BAS population distributions of (A) S-type RuBisCO, (B) F-type RuBisCO, and (C) PepQ aggregates formed in the presence of different amounts of heat-activated IbpAB. Aggregation of RuBisCO-Alexa647 into S-type and F-type aggregates was carried out as outlined in Figure 1. TMR-labeled PepQ was denatured in acid urea and diluted 50-fold to a final monomer concentration of 500 nM in TKM buffer at 50 °C. Aggregation was halted after 4 min by rapid dilution to a final PepQ monomer concentration of 10 nM in TKM buffer at 23°C. The final mixing ratios of heat-activated IbpAB dimer to either RuBisCO or PepQ monomers (1:1, 1:2 and 1:5) are shown. Aggregate particle burst intensity is plotted on the heat map x-axis, while the color scale shows a normalized measurement of particle frequency (fraction of total events observed at each intensity value). Each BAS plot is a combination of n = 3 independent experimental replicates. The number of RuBisCO or PepQ monomers per particle at high IbpAB levels is indicated at the approximate midpoint of the dominant particle population. For these samples, additional dilution (2 to 10-fold) and longer data collection times were employed to permit characterization of the smallest resolvable aggregate particles.

### IbpAB forms distinctive complexes with RuBisCO aggregates

To gain additional insight into some of the properties of these aggregate particles, we examined the size and stoichiometry distributions of the RuBisCO-IbpAB complexes using a two-color variant of BAS (MC-BAS; Figure 3A) (41). MC-BAS employs both the observed raw burst amplitudes and associated burst ratios to reconstruct the size and compositional distributions of two-component nanoparticle systems (41). The same RuBisCO-Alexa647 variant was employed for these experiments. IbpAB was labeled with either Oregon Green on a unique IbpA Cys (D120C) residue, or with an Alexa488 dye on a unique Cys residue added at the C-terminus of IbpB. Neither mutation resulted in a detectable perturbation in aggregation inhibition or aggregate particle size distributions (Figure S3). For these experiments, a 1:1 mixing ratio of RuBisCO monomer to the labeled IbpAB dimer was utilized. While the total labeled particle distributions (i.e. all fluorescent particles, whether with or without detectable RuBisCO monomers) observed with OG-labeled IbpA and Alexa488-labeled IbpB are not identical, the Alexa647 dye on the RuBisCO monomers had little observable impact (Figure S4). At the same time, the Alexa647 dye displays only a small spectral overall with the Oregon Green and Alexa488 dyes, resulting in minimal Förster coupling and little impact on the observed MC-BAS burst distributions (Figure S4 and ref (41)).

**Figure 3.**
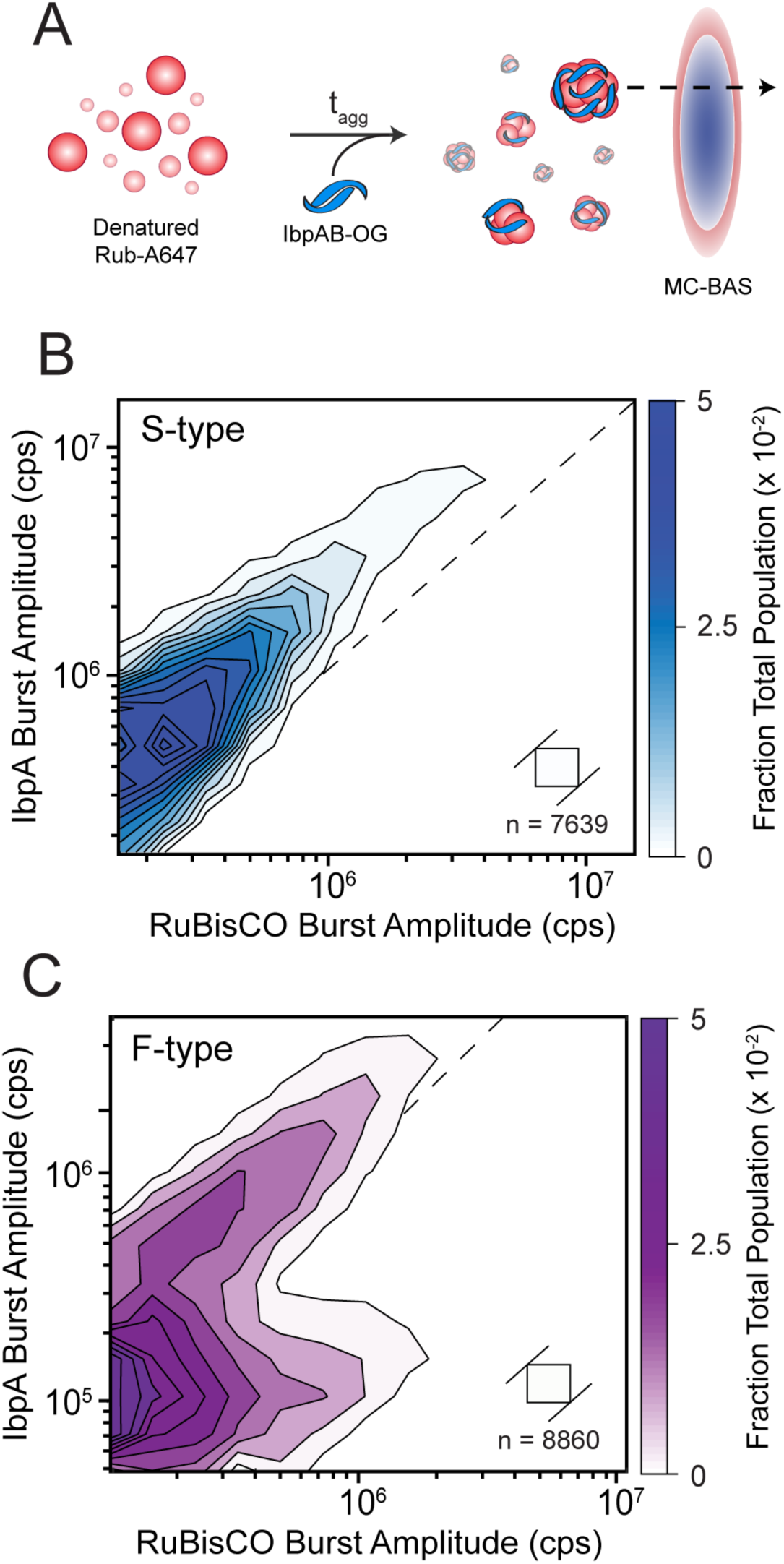
The stoichiometry distributions of IbpAB bound to S-type and F-type RuBisCO aggregates are distinct. (A) S-type and F-type aggregates are formed from RuBisCO-A647 in the presence of heat-activated IbpAB labeled with Oregon Green (IbpA-OG). Observation of co-incident fluorescence bursts in separate detection channels of an MC-BAS microscope demonstrates incorporation of labeled IbpAB into aggregate particles (41). The MC-BAS distributions for (B) S-type and (C) F-type RuBisCO aggregates bound to IbpAB-OG are shown. In each case, the final mixing ratio of RuBisCO monomers to IbpAB dimers was 1:1. The RuBisCO burst intensity is plotted on the x-axis and IbpAB burst intensity is plotted on the y-axis. The dashed diagonal line shows the experimentally determined 1:1 brightness equivalence for the RuBisCO- and IbpA-coupled dyes. The spread of the distributions along the positive diagonals of the plot measures the population size distribution at a given IbpAB:RuBisCO stoichiometry, while the extent of spread along the negative diagonals is proportional to the range of binding stoichiometries. Each MC-BAS plot is a combination of n = 6, independent experimental replicates. The square in each plot shows the 2D bin size prior to contour plot extrapolation and n indicates the total number of coincident burst events in data set.

MC-BAS measurements with labeled IbpA and wild-type IbpB demonstrate that IbpAB forms distinctive co-complexes with similarly sized aggregates (Figure 3B-C). With S-type RuBisCO aggregates, IbpAB binds within a relatively restricted stoichiometry range centered on a RuBisCO:IbpA ratio of approximately 1:2 (Figure 3B). Interestingly, this limited stoichiometry distribution appears to hold across the entire 10-fold aggregate particle size range. However, with F-type aggregates of a similar size, IbpA displays a bifurcated stoichiometry distribution, consistent with the presence of two different sub-populations of IbpAB-RuBisCO particles (Figure 3C). One population is very similar to that observed with amorphous aggregates, displaying a limited stoichiometry distribution centered on a RuBisCO:IbpA ratio of 1:2. The second population appears to incorporate significantly less IbpAB, with the level of IbpA remaining roughly constant even as the aggregate particle size varies by almost 10-fold. Importantly, when the same experiments are conducted with labeled IbpB and wild type IbpA, the same basic patterns and stoichiometry distributions are observed (Figure S5).

### Aggregate disassembly by KJEB is dramatically accelerated by IbpAB

We next examined how different IbpAB-aggregate complexes are disassembled when exposed to the KJEB bi-chaperone disaggregase of *E. coli*. Consistent with our previous observations, S-type aggregates grown at 25 °C for more than a few minutes are only slowly dismantled by the full KJEB system (Figure 4A-C and ref (44)). By contrast, S-type aggregates grown for the same amount of time in the presence of IbpAB are disassembled much more rapidly (Figure 4D-F). Strikingly, particle size appears to have little impact on stimulated disassembly. The size distribution of aggregates formed in the absence of IbpAB largely overlaps the distribution observed when aggregates form in the presence of IbpAB at a 1:1 ratio of IbpAB to RuBisCO (100-2000 RuBisCO monomers; Figure 4B and D). However, particles containing approximately the same number of RuBisCO monomers are only slowly dismantled in the absence of IbpAB (Figure 4B). Near maximal S-type aggregate disassembly is achieved at an IbpAB:RuBisCO ratio of 1:1. While an increased level of IbpAB (up to 5:1) shifts the overall aggregate size distribution more toward smaller particle sizes (100-800 mer), the observed disaggregation rate, estimated from the apparent half-time of particle decay, does not change appreciably (Figure 4E-G). Importantly, MC-BAS experiments with two differently labeled RuBisCO monomer pools demonstrate that no significant re-aggregation takes place during these KJEB-mediated disaggregation experiments (Figure S6 and ref (44)).

**Figure 4.**
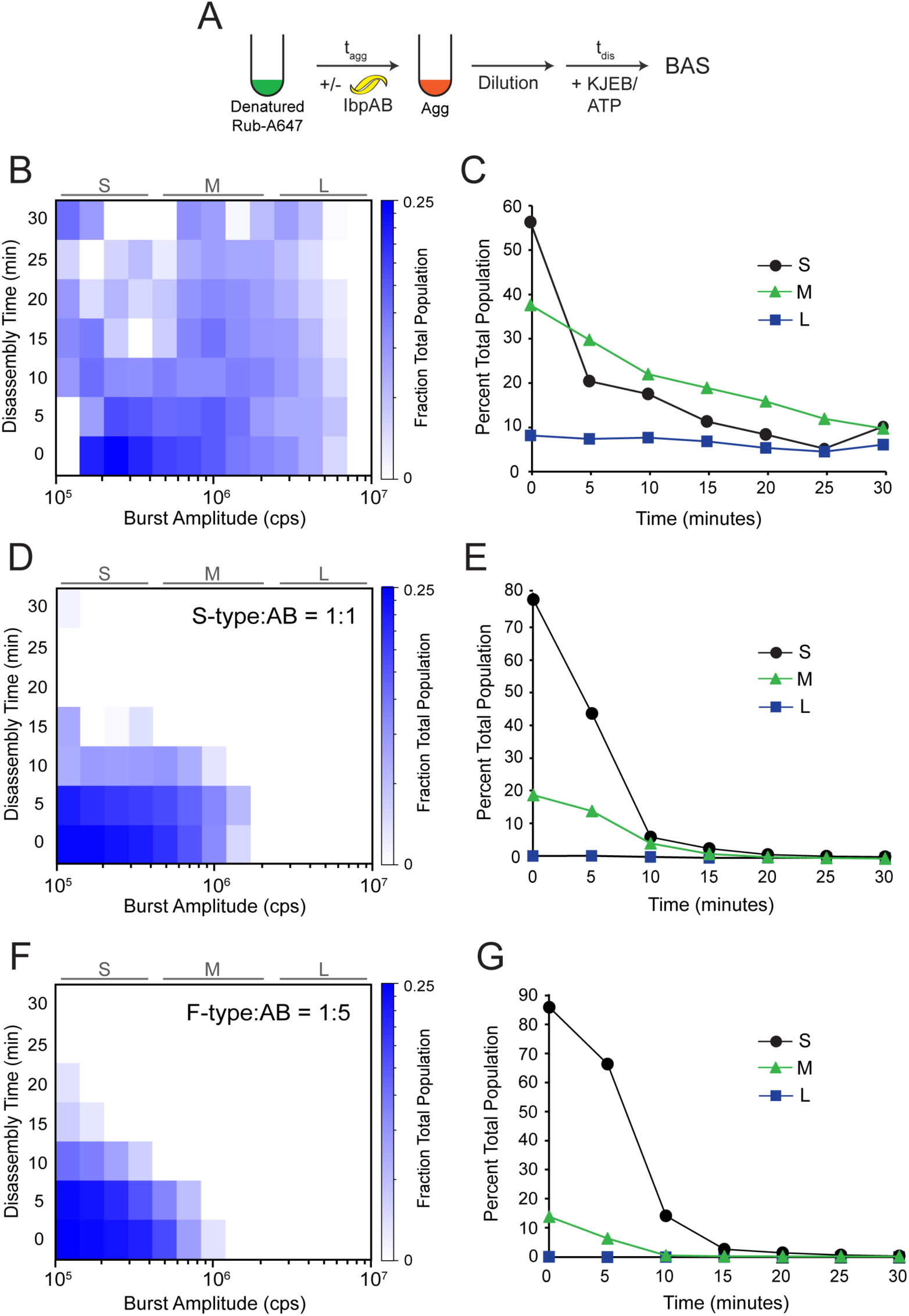
IbpAB dramatically accelerates the disassembly of S-type RuBisCO aggregates by the KJEB bi-chaperone disaggregase. (A) S-type aggregates were formed in either the absence or presence of unlabeled IbpAB. The RuBisCO monomer to IbpAB dimer ratio in each case was: (B) no added IbpAB, (D) 1:1, or (F) 1:5. Disaggregation was triggered by the addition of the KJEB bi-chaperone system (1 µM DnaK, 2 µM DnaJ, 2 µM GrpE and 200 nM ClpB), 2 mM ATP, and a creatine kinase-based ATP regeneration system. Samples were then loaded into a BAS microscope and burst data was continuously acquired for 30 min. The full experimental photon history was segmented into 5 min blocks and analysis was performed on each block. The heat maps represent a combination n = 3, independent experimental replicates for each aggregation condition. A zero-time measurement on each sample was collected prior to the addition of ATP. To highlight how disaggregation rates depended on aggregate size, the BAS heat maps were also coarsely binned into small (*S*), medium (*M*), and large (*L*) particle ranges (C, E, and G). In each case, all detected objects within a given size range were summed and plotted as a function of time following the initiation of disaggregation by KJEB.

IbpAB also dramatically enhances the disassembly of both F-type RuBisCO and PepQ aggregates. F-type RuBisCO aggregates formed in the absence of IbpAB are intrinsically more susceptible to KJEB mediated disassembly than S-type aggregates (Figure 4 B-C and Figure S7 A-B). However, the presence of IbpAB results in a substantial overall increase in aggregate disassembly, though particle size once again has little apparent impact (Figure S7 A-F). Smaller (100-400 mer) and medium sized (500-1000 mer) F-type aggregates display distinctive disassembly half times in the absence of IbpAB (Figure S7 B). However, aggregates formed in the presence of IbpAB that have these same number of RuBisCO monomers, display disassembly half times that are all similar (Figure S7 D-F). By contrast, PepQ aggregates are completely resistant to disassembly by KJEB in the absence of IbpAB (Figure S8 A-B). Interestingly, when IbpAB is present with PepQ at a 1:1 ratio, the aggregate particle distribution displays a large shift toward smaller particles (Figure 2C and Figure S8C). However, only the smallest particles at this IbpAB:PepQ ratio show any, and only very limited, susceptibility to disassembly, with medium sized particles remaining almost totally resistant (Figure S8 C-D). Increasing the IbpAB level to a 5:1 ratio results in a complete shift of the aggregates into a smaller particle size range and a dramatic enhancement in disassembly (Figure S8 E-F). Once again, particle size does not predict disaggregation potential. Small PepQ aggregates that form in the absence of IbpAB, and which exist in the same size range as the IbpAB-induced particles, are highly resistant to disassembly (Figure S8 A-B).

### Incorporation of IbpAB results in expansion of aggregate structure

We next asked whether the co-assembly with IbpAB alters the structure of RuBisCO aggregates. For these experiments, we employed a set of previously described *inter*- and *intra-*molecular FRET assays that report on the average conformational properties of RuBisCO monomers and aggregates (42–44, 47, 48). For *inter-*molecular FRET measurements, different pools of RuBisCO monomer carry the donor or acceptor probe. Upon co-aggregation, Förster coupling reports on the relative proximity of the labeled segments of the RuBisCO monomer averaged across the total aggregate population (Figure 5A; (43, 44). In the case of *intra*-molecular assays, the same RuBisCO monomer carries both donor and acceptor probes (Figure 5D). When the double-labeled monomer is co-aggregated with a much larger pool of unlabeled monomer, the observed FRET signal primarily reports on the average conformation of individual, aggregate-incorporated monomers (42–44, 47, 48).

**Figure 5.**
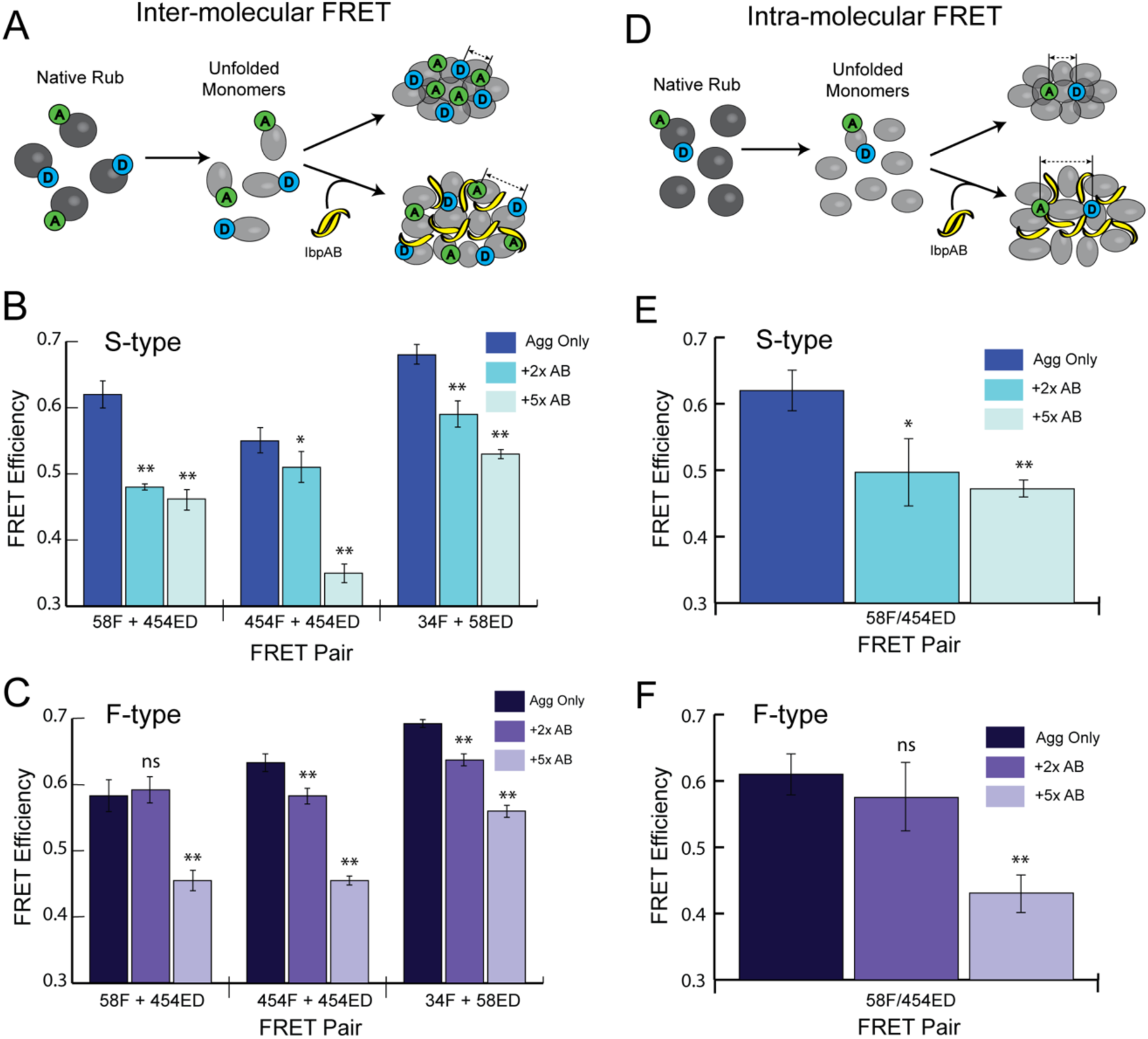
IbpAB induces expansion of both aggregate particle structure and aggregate-incorporated monomer structure. (A) Schematic of an experiment using ensemble intermolecular FRET to examine the impact of IbpAB on the average relative proximity of RuBisCO monomers within aggregates. S-type (B) and F-type (C) aggregates were prepared either in the absence or presence of excess IbpAB. Samples of RuBisCO, either unlabeled, labeled with a donor fluorophore only (IAEDANS) or labeled with an acceptor fluorophore only (fluorescein) were denatured and pair-wise mixed at 1:1 ratio prior to initiation of aggregation in the presence or absence of either a 2-fold or 5-fold excess of IbpAB. Three concentration-matched samples were thus created for each aggregate type, both with and without IbpAB: donor-only, acceptor-only or donor-acceptor. The same protocol was replicated using three different paired labeling sites on the RuBisCO monomer (58F + 454ED; 454F + 454ED; 34F + 58ED; (42, 43)). FRET efficiencies are calculated from the magnitude of the corrected donor-side quenching, while enhanced acceptor fluorescence (not shown) was used to confirm Förster coupling. (D) Schematic of an experiment using ensemble intramolecular FRET to examine the impact of IbpAB on the average conformation of aggregate-incorporated RuBisCO monomers. S-type (E) and F-type (F) RuBisCO aggregates were created by mixing a denatured, doubly labeled RuBisCO monomer (58F/454ED; (43, 44)) into a large excess of denatured, unlabeled RuBisCO (1:9), prior to the initiation of aggregation, in the presence or absence of a 2-fold or 5-fold excess of IbpAB. Under these conditions, Förster coupling between different labeled monomers within the same aggregate particle is minimal, so that the observed FRET signal is dominated by coupling between the probes attached to the same monomer. Concentration-matched reference samples using donor-only (454ED) and acceptor-only (58F) RuBisCO were also prepared and used to calculate the donor-side FRET efficiency, as well as confirm the presence of Förster coupling. In all cases, error bars show the s.d. of n = 3 independent experimental replicates. The differences in FRET efficiency of the aggregate-only samples compared to those containing IbpAB were evaluated for statistical significance using a two-tailed student’s t-test (ns, not significant; *, P < 0.005; **, P < 10^-5^).

The impact of IbpAB on both S-type and F-type RuBisCO aggregates is consistent with a generalized expansion of aggregate structure. Using three distinct FRET pairs, we first examined the relative proximity of different segments of the RuBisCO monomer within aggregates. In all cases, and for both S-type and F-type aggregates, the addition of IbpAB results in a reduction in average FRET efficiency (Figure 5 B and C). Next, we examined the average conformation of an aggregate-incorporated monomer along one principal axis between the N- and C-terminal domains (43). With both S-type and F-type aggregates, incorporation of IbpAB results in an increase in the average distance between the N- and C-terminal regions of the aggregate-incorporated RuBisCO monomer (Figure 5 E and F). Interestingly, with F-type aggregates, higher levels of IbpAB (5:1) are required to achieve the maximally observed change in FRET efficiency for both inter- and intra-molecular measurements. While some of the observed reduction in FRET efficiency could result from reductions in overall particle size (Figure 2), the magnitude of this effect is likely small. Using the same assay, we previously showed that the average particle FRET efficiency reaches a maximum at the earliest stages of particle growth and becomes rapidly insensitive to size (44). Consequently, at the aggregation time points used here, the observed FRET efficiency is dominated by the relative proximity of the donor and acceptor probes within assembled particles rather than the overall number particles.

### IbpAB release and aggregate particle disassembly are coincident

To gain additional insight into the process of stimulated disaggregation, we used MC-BAS to observe, in real time, aggregate disassembly and IbpAB release (Figure 6A). We employed the same labeled RuBisCO and IbpAB outlined in Figure 3. In this case, the RuBisCO:IbpAB mixing ratio was again set at 1:1 and the concentration of the KJEB system was reduced in order to slow disassembly and improve event detection and temporal resolution during disaggregation.

**Figure 6.**
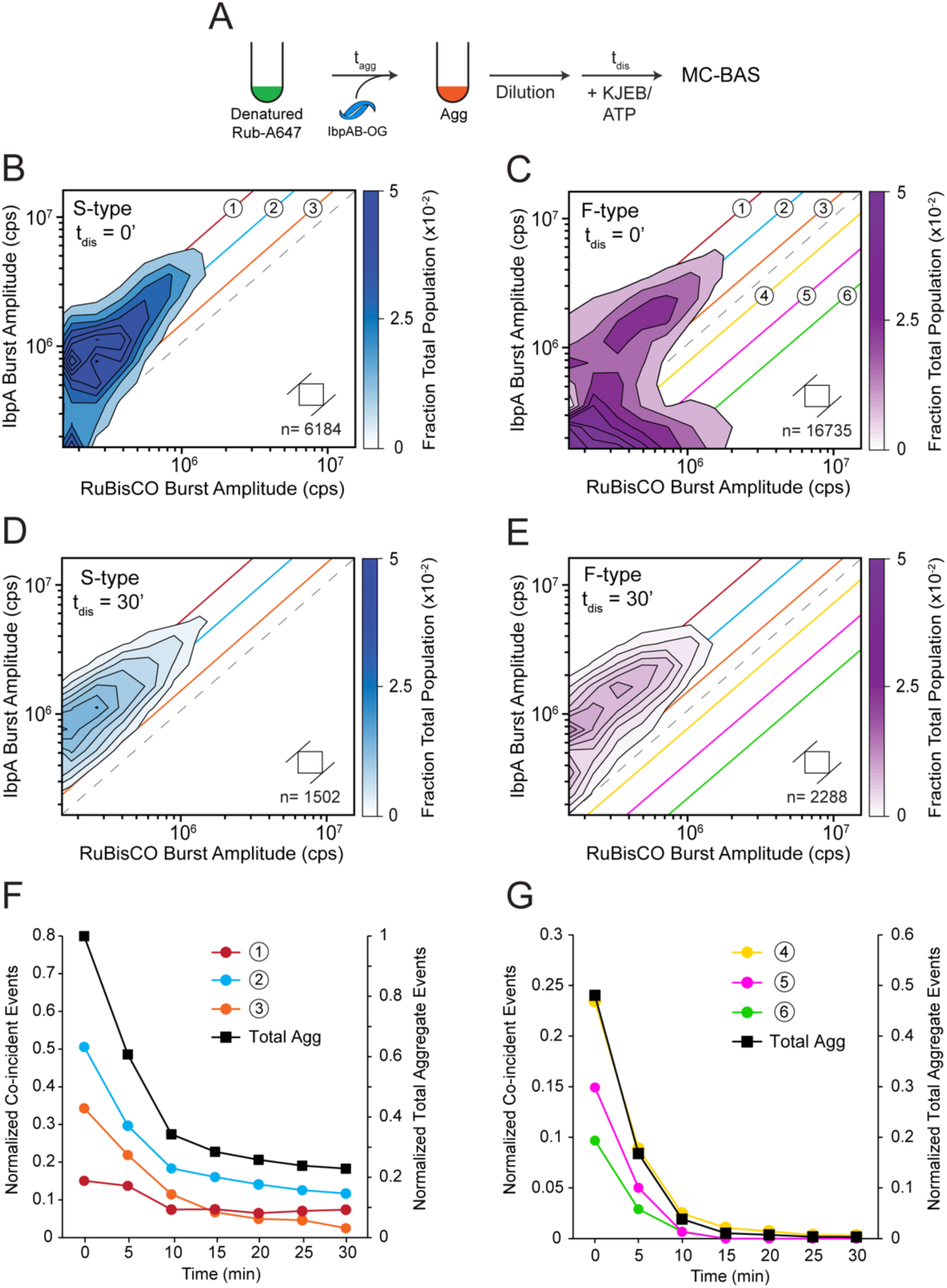
IbpAB removal from RuBisCO aggregates closely tracks overall particle disassembly. (A) Aggregates were created from RuBisCO-A647 in the presence of IbpA-OG and wild-type IbpB at a 1:1 ratio of RuBisCO monomer to IbpAB dimer. Disaggregation was triggered by addition of the KJEB bi-chaperone system (250 nM DnaK, 500 nM DnaJ, 500 nM GrpE and 50 nM ClpB), 2 mM ATP, and a creatine kinase-based ATP regeneration system. Samples were then loaded onto a BAS microscope and burst data was continuously acquired for 30 min. Each MC-BAS plot is a combination of n = 6, independent experimental replicates. (B and C) A zero-time MC-BAS measurement was collected for both S-type and F-type aggregates prior to the addition of ATP. The experimentally calibrated 1:1 stoichiometry ratio is indicated by the gray dashed line. Colored diagonals illustrate coarse binning ranges for the IbpAB:RuBisCO binding stoichiometry at both higher (*1, 2* and *3*) and lower (*4, 5* and *6*) ratios. After 30 min of KJEB disassembly, both S-type (D) and F-type (E) aggregates show similar residual co-labeled particle distributions. (F and G) The kinetics of IbpAB release from aggregates in each coarse stoichiometry range, observed as the disappearance of co-labeled particles, relative to the rate at which the same aggregate particles are disassembled (*Total Agg*), is shown. For S-type aggregates (F) only the higher IbpAB:RuBisCO stoichiometry ranges are significantly populated. For F-type aggregates, co-labeled particles in both higher and lower IbpAB:RuBisCO stoichiometry ranges are observed. The kinetics of IbpAB release from the higher stoichiometry sub-population of fibril-like aggregates (C; *1, 2* and *3*) is similar to the behavior of amorphous aggregates. The release of IbpAB from the lower stoichiometry sub-population of the F-type aggregates, relative to the rate at which these particles are disassembled, is shown in (G). The reduced normalized amplitude observed in panel G is a consequence of the particles observed in the lower IbpAB:RuBisCO stoichiometry range (4–6) representing approximately half of the total observed particle population (C). The square in each plot shows the 2D bin size prior to contour plot extrapolation and n indicates the total number of coincident burst events in data set.

Addition of KJEB and ATP to either IbpAB-bound S-type or F-type aggregates triggers a time-dependent disappearance of coincident objects across the full range of particle sizes (Figure 6B-E). The co-labeled S-type aggregates again initially cluster around a binding stoichiometry of 1:2 (Figure 3) and, surprisingly, display no detectable shift in stoichiometry distribution as they decay (Figure 6B and D). Coarse binning of the amorphous aggregate particles by their RuBisCO:IbpAB stoichiometry shows that particles containing both higher and lower levels of IbpAB all disappear at essentially the same rate (Figure 6A, D and F). The decay of these coincident particles closely tracks the rate at which the S-type RuBisCO aggregates themselves are dismantled (Figure 6F). When fluorescence bursts are separated by strict channel coincidence, a distinct sub-population of S-type aggregate particles is detected (Figure S9). This sub-population either does not bind IbpAB or does so at such a low level that the bound IbpAB cannot be detected. These IbpAB-depleted particles, nonetheless, remain highly susceptible to disassembly (Figure S9 E). At the same time, a sub-population of particles with a similar number of RuBisCO monomers, but which is much more enriched in IbpAB, appear to be more slowly disassembled (Figure S9 C-D). Non-coincident IbpAB events, which appear unbound to any RuBisCO subunits, accumulate over time, consistent with the release of IbpAB from aggregates and their accumulation in larger oligomers with 50-400 subnits (Figure S9 F).

Despite their more complex subpopulation composition, similar results are obtained with the F-type aggregates (Figure 6C-G). The particle subpopulation that is enriched in bound IbpAB, and which has a stoichiometry distribution similar to the S-type aggregates (Figure 3 and Figure 6B and C), follows a coincidence decay pattern that is very similar to that observed with the amorphous aggregates (data not shown). Likewise, the F-type subpopulation that is depleted in IbpAB disappears without any apparent dependence on particle size and with no detectable shift in stoichiometry distribution (Figure 6C and E). Coarse binning of this sub-population by stoichiometry ratio demonstrates that, while these particles are dismantled more rapidly and more fully than the other subpopulation or the S-type aggregates, their decay is also independent of IbpAB level and, once again, closely tracks the overall disassembly of the RuBisCO particles themselves (Figure 6G). Thus, release of IbpAB from both S-type and F-type aggregate co-complexes occurs at the same overall rate at which RuBisCO monomers themselves are disassembled by the KJEB bi-chaperone system.

### Stimulated disaggregation requires incorporation of IbpAB early in aggregate growth

We next examined whether stimulated disaggregation requires that IbpAB be present at the earliest stages of aggregation. To address this question, we modified the co-assembly protocol so that activated IbpAB can be added at different delay times relative to the onset of RuBisCO aggregation (Figure 7A). We then used both BAS and MC-BAS to determine whether delayed IbpAB addition arrests aggregate growth while also characterizing the stoichiometry distribution of any resulting co-complexes.

**Figure 7.**
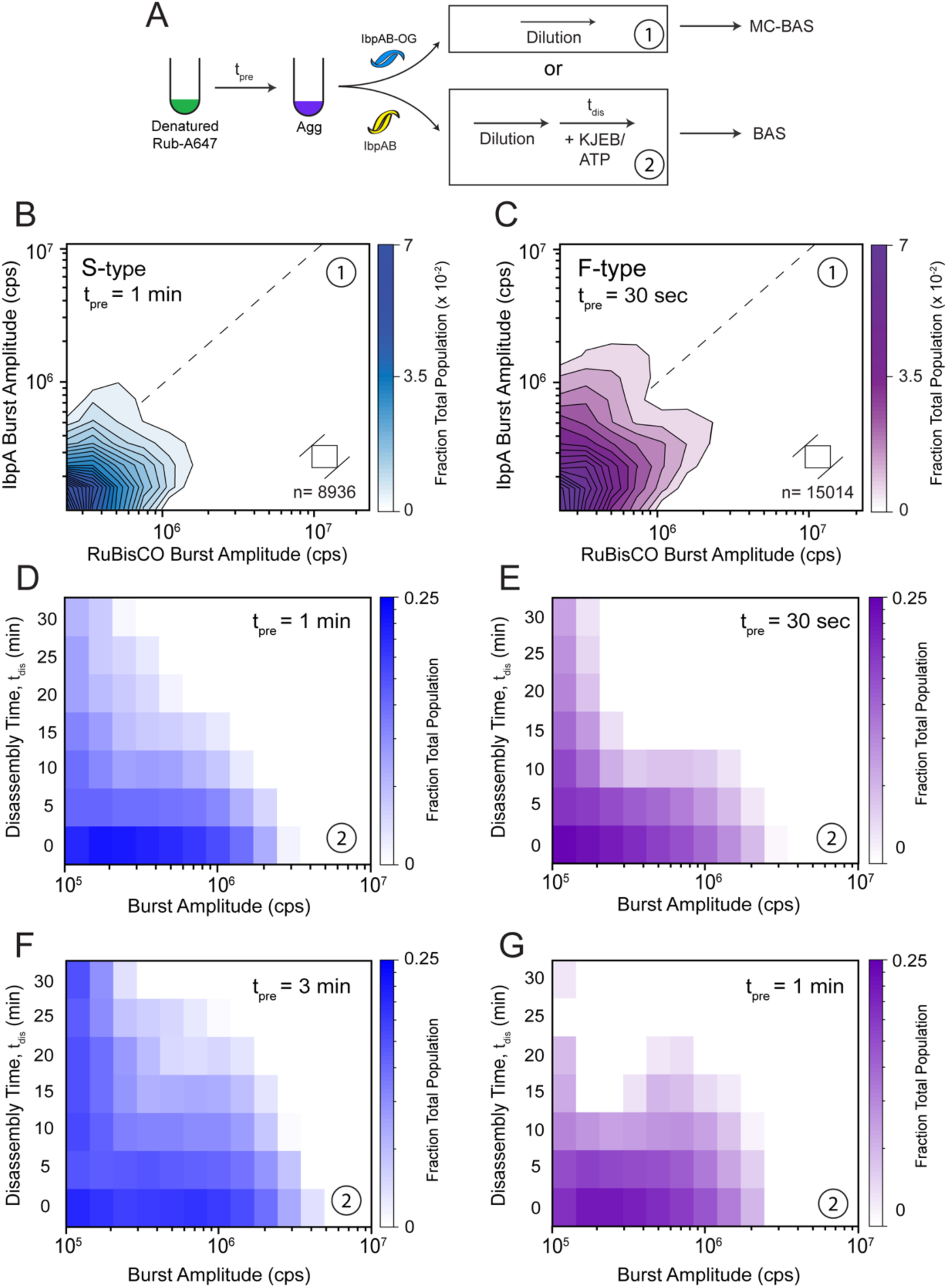
Delayed addition of IbpAB to RuBisCO aggregates prevents further particle growth but results in loss of stimulated disassembly. (A) Sample handling protocol for experiments where the addition of IbpAB to an aggregating sample of RuBisCO is delayed (*t_pre_*) relative to the initiation of aggregation. MC-BAS was used to examine the binding distribution of IbpAB-OG on S-type (B) and F-type (C) RuBisCO-A647 aggregates upon delayed IbpAB addition in the absence of KJEB-mediated disassembly. For S-type aggregates, t_pre_ = 1 min and for F-type aggregates, t_pre_ = 30 sec. In both cases, the final RuBisCO monomer:IbpAB mixing ratio was 1:1 and no KJEB disaggregase was added. Each MC-BAS plot is a combination of n = 6, independent experimental replicates. The population-resolved kinetics of disaggregation by KJEB were examined with BAS following delayed addition of wild type IbpAB to S-type (D and F) and F-type (E and G) RuBisCO aggregates. For S-type aggregates, t_pre_ = 1 min (D) or 3 min (F) at a final RuBisCO monomer to IbpAB mixing ratio of 1:1, while for the F-type aggregates a final mixing ratio of 1:5 and t_pre_ = of 30 sec (E) and 1 min (G) were used. Disassembly conditions and KJEB concentrations were identical to those used in Figure 4. Each heat map is a combination of n = 3 independent experimental replicates.

For both S-type and F-type RuBisCO aggregates, delayed addition of IbpAB at a 1:1 mixing stoichiometry halts further aggregate growth and produces a substantial population of co-labeled particles (Figure 7B-C and Figure S10). However, the resulting stoichiometry distributions are distinct from those observed when IbpAB is added at the beginning of aggregation. In both cases, co-labeled particles are mainly populated at the small end of the aggregate size range (Figure 7B-C). Additionally, while the F-type aggregates appear to form a small population of particles that exceed a RuBisCO:IbpAB ratio of 1:1, this population is almost completely absent with S-type aggregates, which incorporate much less IbpAB when their addition is delayed (Figure 7B). The previous bifurcation of the F-type particles into two, co-labeled subpopulations is also far less distinct (Figure 7C).

Delayed addition of IbpAB also reduces enhanced disaggregation of both aggregate types. Delaying the addition of IbpAB to S-type aggregates by 1 min, relative to the onset of aggregation, results in a particle size distribution that is only slightly larger than that observed when IbpAB is added at the beginning of aggregate growth (Figure 7D and Figure S10). However, the rate of aggregate disassembly slows substantially over the entire particle size range, with a subpopulation of smaller particles persisting even after 30 min of disassembly (Figure 7D). When the delay is increased to 3 min, disaggregation slows more, even though the aggregate particles mostly populate a similar overall size range (Figure 7F). Interestingly, delayed addition of IbpAB to F-type aggregates does not result in as dramatic a loss of enhanced disassembly (Figure 7E and G). When addition of IbpAB at a mixing ratio of 1:5 is delayed by up to 1 min, disassembly of F-type aggregates slows compared to a similar addition of IbpAB at the beginning of aggregate growth (Figure 7G). Nonetheless, F-type aggregates are still dismantled more effectively than are S-type aggregates (Figure 7F and G). At the same time, delayed addition of IbpAB at a mixing ratio of 1:5 results in disaggregation kinetics that are similar to the un-delayed addition of IbpAB at a mixing ratio of 1:1 (Figure 7G and FigureS7 C and D). Overall, these observations suggest that optimal enhancement of disaggregation by IbpAB requires their incorporation at the earliest stages of aggregate particle growth.

## DISCUSSION

We previously showed that the correlation between RuBisCO aggregate particle size and the observed rate of KJEB disassembly is weak, suggesting that other features of an aggregate particle are much more important in determining the efficiency and rate of this key protein quality control process (41). However, the generality of these observations was limited by: (1) the characterization of a single substrate protein (2) the heterologous source of the substrate protein (*R. ruburm*) and disaggregase chaperones (*E. coli*) and (3) the absence of small heat shock chaperones, which would normally be present at the earliest stages of protein aggregation *in vivo*. The examination of PepQ aggregation and disaggregation, in the presence and absence of IbpAB, addresses these issues (Figure 2 and S8). PepQ, like RuBisCO, displays stringent refolding behavior that requires the action of the GroELS chaperonin system to efficiently form a natively folded enzyme. At the same time, PepQ has a distinctive amino acid sequence, a fundamentally different native structure, and significantly different aggregation behavior (45, 46). But, as an endogenous *E. coli* enzyme, PepQ permits evaluation of whether specific coupling exists between an sHsp system and a co-evolved substrate protein. The very similar behavior we observe with RuBisCO and PepQ argues strongly that the behavior we observe is likely to be general.

The dramatic stimulation of RuBisCO and PepQ disaggregation in the presence of IbpAB suggests that co-complexes formed between these sHsp chaperones and their client proteins are, in some way, optimized for efficient disassembly. While our results are consistent with prior studies of other model proteins, where enhanced disaggregation is seen in the presence of sHsps, they are not consistent with some mechanisms proposed to explain this enhanced disaggregation. For example, enhanced disaggregation in the presence of sHsps is typically associated with suppressed aggregation and reduced average aggregate particle size (13–15). This correlation has been interpreted to indicate that aggregate size is a primary limiting physical property on disassembly. However, our observations demonstrate that the number of RuBisCO or PepQ monomers incorporated into an aggregate particle, either with or without IbpAB, does not strongly correlate with disassembly rate (Figure 4 and S7-S8). It has also been suggested that sHsps like IbpAB form shells around the exterior of a growing aggregate particle in order to block exposed interaction surfaces of the client proteins necessary for aggregate particle growth, limiting aggregate size to more manageable dimensions (27). In principle, disassembly of such an encapsulated aggregate particle could be expected to display a burst of IbpAB release that precedes aggregate particle disassembly, as the KJEB disaggregase system first dismantles the outer sHsp shell (27). However, our two-color MC-BAS experiments detect no burst of IbpAB release that precedes aggregate disassembly (Figure 6). Within the temporal resolution of our experiments, IbpAB release from both RuBisCO and PepQ aggregate particles appears to be coincident with client protein disassembly.

Our observations are also not consistent with models in which sHsps act to maintain client proteins in near-native conformations to facilitate subsequent reactivation during disaggregation, as has been proposed for some eukaryotic sHsps (30–32). Neither RuBisCO nor PepQ refold to any detectable extent upon release from IbpAB particles and remain fully dependent on GroELS. Complete folding of these stringent GroEL substrate proteins is not necessary for efficient disaggregation, either in the presence or absence of IbpAB, so that rapid disaggregation and efficient folding cannot be obligately coupled. At the same time, disassembly of RuBisCO aggregates, both in the presence and absence of IbpAB does not lead to detectable re-aggregation, so long as the KJE system remains active (Figure S6).

We hypothesize that IbpAB follows an adaptive assembly mechanism when forming co-particles with non-native client proteins (Figure 8). In such a model, the amount of IbpAB required to create a limit aggregate particle is coupled to properties of the aggregating protein, while the target size of the final aggregate is encoded in the assembly behavior of IbpAB itself. Consistent with this hypothesis, IbpAB is capable of restricting three different aggregate types to a similar number of monomers per particle, despite their widely varying physical properties. At the same time, different levels of IbAB are required to form these limit aggregate particles and stimulate their disaggregation. For example, while IbpAB shifts F-type RuBisCO aggregates toward smaller particle sizes, it appears to only modestly enhance the overall disassembly of this sized aggregate. However, even at an IbpAB:RuBisCO ratio of 1:1, medium sized particles are disassembled much more rapidly than in the absence of IbpAB (Figure S7 A-D). When the IbpAB ratio is increased to 5:1, the residual medium sized particles disappear even more quickly (Figure S7 E-F). MC-BAS analysis is also consistent with this hypothesis (Figure 3 and S5). These experiments show that IbpAB can be incorporated into sub-populations of F-type RuBisCO aggregates at distinct binding stoichiometries, despite these particles possessing similar numbers of non-native RuBisCO subunits and disassembling at similar overall rates.

**Figure 8.**
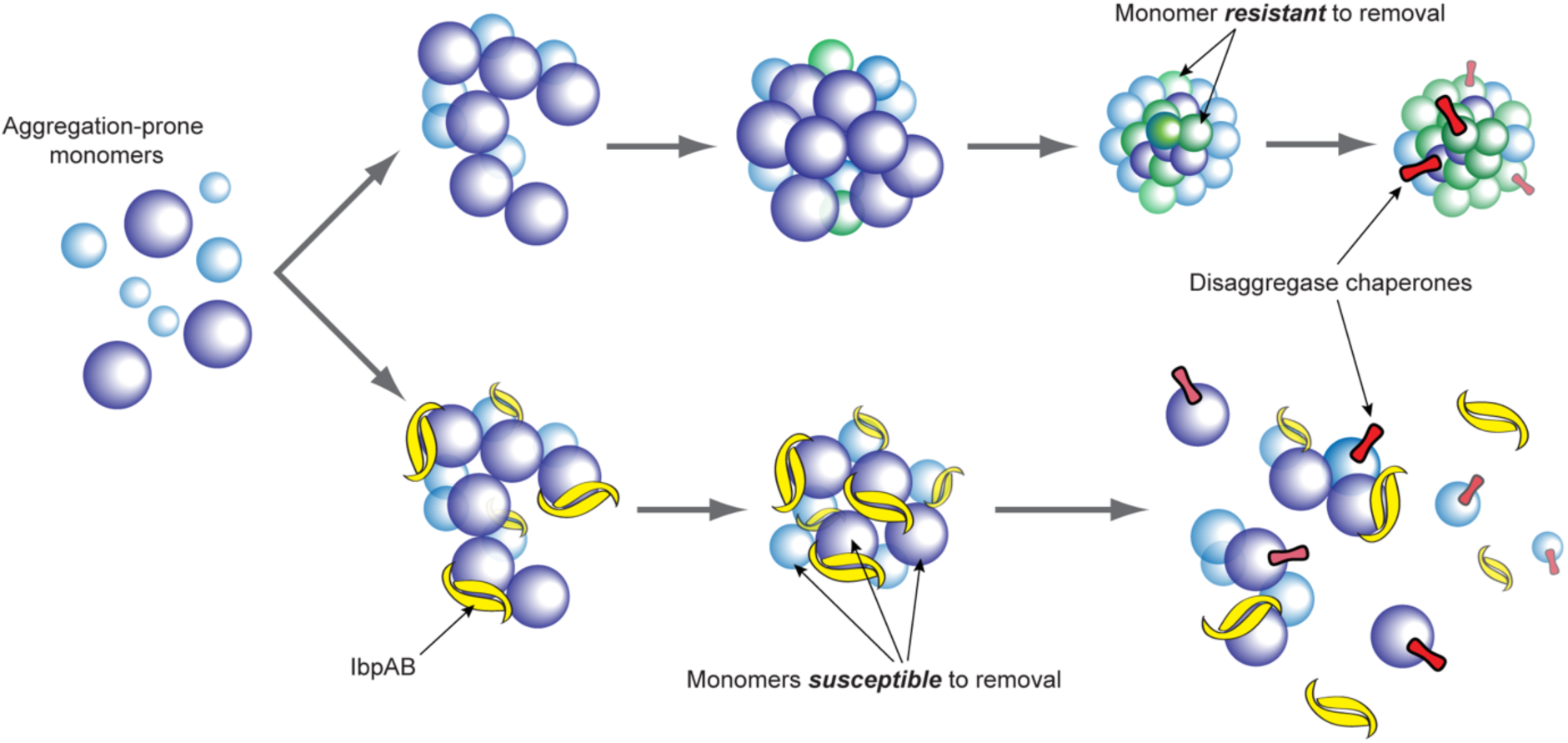
Proposed model for IbpAB-enhanced protein disaggregation via internal structural modification of aggregate particles. Aggregate formation in the absence of IbpAB (upper) results in particles that rapidly change subunit conformation or packing interfaces, or both, so that the energy needed to remove a monomer exceeds that available to the bi-chaperone disaggregase system. In the presence of IbpAB (lower), subunit conformational changes and/or packing interfaces are maintained in a configuration that is more readily disrupted by the bi-chaperone disaggregase. IbpAB thus acts as a “lubricant” to facilitate extraction or sliding of aggregate subunits against one another during disassembly by the ATP-powered bi-chaperone disaggregase.

In total, our results are most consistent with a model of aggregate disassembly in which structural and/or dynamic features of the aggregated client subunits, along with their packing interactions, dominate whether or not an aggregate particle can be dismantled (Figure 8). We have shown, here and in previous work (44), that the KJEB bi-chaperone disaggregase can disassemble both F-type and S-type RuBisCO aggregates in the absence of IbpAB. However, this activity is maximal at relatively early time points in the aggregate growth process, with aggregates becoming progressively resistant to disassembly as they age (44), a phenomenon that is not reversed by late addition of IbpAB (Figure 7). Previous FRET measurements suggested that changes in internal aggregate structure, changes consistent with a time-dependent increase in the compaction of both individual monomers, as well as subunit packing interfaces, are a better predictor of RuBisCO aggregate susceptibility to KJEB alone (44). Similar FRET analysis in the presence of IbpAB further supports these conclusions (Figure 5) and are consistent with observations using other model proteins. Our observations support a model of IbpAB action in which direct incorporation of IbpAB into, and modification of, the structure of aggregate particles is the essential mechanism of action for this sHsp system (Figure 8) (28, 29). Our observations further suggest that IbpAB may be capable of sensing general features of nascent aggregate formation and guiding very different proteins along similar co-assembly pathways that are maximally amenable to subsequent disassembly. It is tempting to speculate that IbpAB acts as a general “lubricant” within co-aggregate particles, facilitating the removal, extraction or sliding of subunits against one another (Figure 8). The nature of such a general lubricating property, if it exists, remains unclear and will require additional investigation.

## METHODS

### Protein Expression and Purification

Wild-type RuBisCO was expressed and purified as previously described (42, 43). Labeling of the single, surface exposed Cys residue (Cys58) with either Alexa488-maleimide or Alexa647-maleimide (ThermoFisher) was conducted as previously described (42, 43, 49). In each case, the labeling efficiency (> 98%) was determined as previously described (50, 51). PepQ A24C was expressed, purified and labeled with TMR-5-iodoacetamide dihydroiodide (Invitrogen) as previously described (45), with a labeling efficiency of > 95%. DnaK, DnaJ, GrpE and ClpB were expressed, purified and stored as previously described (44).

Both wild-type and mutants of IbpA and IbpB were expressed and purified as previously described (34–36), with some modifications. In brief, IbpA and IbpB were individually expressed as N-term poly-His fusion proteins in *E. coli* BL21(DE3) at 37°C using 400 uM IPTG for 3 hrs. Following high-pressure shear lysis (M110Y, Microfluics International) and clarification by centrifugation (95,000x g), crude lysates were loaded onto a Ni-NTA Column (Qiagen) under denaturing conditions in urea buffer (50 mM Tris pH 7.4, 300 mM NaCl, 20 mM imidazole, 6 M deionized urea, 5 mM BME). Fusion proteins were eluted from the column with a single 500 mM imidazole step, and fractions containing IbpA or IbpB were collected and combined. Proteins were renatured using a step-wise reduction in urea concentration by dialysis (4 M, 2 M, 1 M urea in urea buffer) at 4 °C, followed by multiple additional exchanges in urea buffer minus the urea to remove residual urea. The poly-His affinity tag was then removed by incubation with TEV protease at 4°C for 24 hours. Samples of IbpA or IbpB were again denatured in urea buffer and loaded onto a Ni-NTA affinity column to remove TEV protease and any un-cleaved fusion protein. Fractions containing IbpA and IbpB were collected, combined and loaded onto a strong anion exchange column (MonoQ, GE) in denaturing MonoQ buffer (50 mM Tris pH 7.4, 6 M deionized urea, 2 mM DTT) and eluted with a linear NaCl gradient (0 - 1 M) over 25 CV. Final renaturation of either IbpA or IbpB was accomplished using a stepwise reduction in the urea concentration by dialysis as described above into IbpAB storage buffer (25 mM Tris pH: 7.4, 150 mM KCl, 0.5 mM EDTA, 2 mM DTT), supplemented with glycerol, (15% v/v) and snap frozen using liquid N_2_.

### Protein Labeling

Mutants of IbpA (D120C) and IbpB (143C) were labeled with the fluorescent, thiol-reactive dyes Oregon Green-maleimide (OG) and Alexa-488-maleimide (ThermoFisher). Each reactive dye was freshly prepared from dry powder in anhydrous DMF immediately prior to use. Labeling was accomplished using a protocol similar to that previously described for RuBisCO and PepQ (42, 43, 45, 49), with some modifications. Both native IbpA (D120C) and IbpB (143C) were separately reduced with 1 mM TCEP in labeling buffer (50 mM Tris pH 7.4, 150 mM KCl, 0.5 mM EDTA) through overnight dialysis at 4°C. Following dye addition and incubation at 23 °C, the reaction was quenched with glutathione, followed by addition of buffered, deionized urea to a final concentration of 3 M. Unreacted dye was removed by sequential dilution and concentration, followed by gel filtration (PD-10, GE) in the same urea buffer. Labeled protein samples were then rapidly diluted to 500 mM urea and dialyzed against IbpAB storage buffer overnight at 4 °C to fully remove residual urea. The protein was then concentrated in the same dialysis tubing using a dry bed of PEG 20,000, collected, dialyzed into fresh storage buffer, supplemented with glycerol (15% v/v), and finally snap frozen in liquid N_2_.

### Protein Aggregation

Native RuBisCO was first denatured in acid urea buffer (25 mM glycine-phosphate, pH 2.0, 8 M deionized urea) at 10 μM for 30 min at 23°C. Denatured RuBisCO was then rapidly diluted (50-fold; 200 nM final monomer concentration) into HKM buffer (50 mM HEPES, pH 7.6, 150 mM KOAc, 10 mM Mg(OAC)_2_, 2 mM DTT) at either 23°C for 2 minutes (F-type) or for 2 minutes at 4°C, followed by incubation at 23°C for 5 minutes (S-type). In both cases, the initial dilution (300 μl total volume) was carried out using a 1 ml non-stick centrifuge tube with a magnetic spin vane placed on a stir plate set for maximum stirring velocity (∼ 1500 rpm). Stirring was halted after 10 sec. Following final incubations, aggregation was halted by dilution (20-fold) into HKM buffer at 23 °C to a final monomer concentration of 10 nM monomer (44).

Native PepQ was denatured in acid urea buffer at 10 μM for 30 min at 23°C. Denatured PepQ was then rapidly diluted (20-fold; 500 nM final monomer concentration) into TKM buffer (50 mM Tris pH, 7.4, 50 mM KOAc, 10 mM Mg(OAc)_2_, 2 mM DTT) at 50°C using the same non-stick centrifuge tube method described above. The sample was incubated at 50°C for an additional 4 minutes, followed by dilution (20-fold) to a final PepQ monomer concentration of 10 nM to halt aggregation.

Before use in aggregation experiments, mixtures of IbpA and IbpB (1:1) were heat activated at 42°C for 10 minutes in HKM or TKM buffer. Samples of activated IbpAB were then added to either HKM buffer (RuBisCO) or TKM buffer (PepQ) immediately prior to addition of denatured RuBisCO or PepQ. The IbpAB dimer concentration (10 nM - 50 nM) was varied in different experiments, relative to the RuBisCO or PepQ monomer concentration, as indicated.

### Protein Disaggregation

In all cases, disaggregation was conducted in a reaction volume of 1 ml, at a final RuBisCO or PepQ monomer concentration of 10 nM. The aggregate sample was supplemented with the bi-chaperone disaggregase from *E. coli*: DnaK (1 µM), DnaJ (2 µM), GrpE (2 µM), and ClpB (200 nM). Where indicated, the concentration of the bi-chaperone disaggregase was reduced, but the relative component ratios remained the same. In all cases, disaggregation was triggered by the addition of ATP (2 mM) and a creatine phosphate/creatine kinase ATP regeneration system (44).

### Förster Resonance Energy Transfer (FRET) Experiments

Aggregation of RuBisCO was examined by FRET using a set of previously described assays (43, 44). For inter-molecular FRET, two different RuBisCO samples were employed, each of which carried either a donor (AEDANS) or acceptor (fluorescein) probe. For intra-molecular FRET, the donor and acceptor dyes were site-specifically conjugated to the same RuBisCO monomer at position 58 (fluorescein) or 454 (AEDANS) (43). For all inter-molecular FRET experiments, 200 nM of both the donor and acceptor labeled monomers (400 nM total) were mixed in acid urea prior to the initiation of aggregation. For intra-molecular FRET, the doubly labeled RuBisCO monomer was mixed in acid urea with an excess of unlabeled wild-type RuBisCO (400 nM final total monomer, 75-90% unlabeled), in order to minimize Förster coupling between different doubly labeled monomers within the same aggregate particles. Following the same protocols described above, either amorphous or fibril-like aggregates were formed in the presence or absence of IbpAB. Aggregation was halted by a 40-fold dilution of the aggregating sample to a final RuBisCO monomer concentration of 10 nM prior to fluorescence measurement. Each FRET measurement involved three, concentration matched samples: (1) donor-only, (2) donor-plus-acceptor and (3) acceptor-only. Identical aggregation conditions and concentrations were employed for all samples in an experimental set, with the donor-only and acceptor-only samples employing the relevant single labeled RuBisCO monomers and wild-type, unlabeled monomers.

Steady state fluorescence emission spectra were acquired with a T-format photon-counting spectrofluorometer (PTI) equipped with a temperature-jacketed cuvette holder maintained at 23 °C. Sample excitation was set at 336 ± 6 nm and emission was collected from 420-550 ± 6 nm. The observed, average FRET efficiency (E) was calculated from the background-corrected donor emission spectra in the presence (FDA) and absence (FD) of the acceptor, by integration of the donor emission over wavelengths where no acceptor emission was detectable: E = (FD - FDA) / FD. The observed acceptor-side FRET signal was used to qualitatively confirm the existence of energy transfer.

### BAS Data Collection and Analysis

Fluorescence burst data was acquired with a previously described custom-built, multi-channel BAS microscope (40, 41). Following initiation of disaggregation by the addition of ATP, samples (10 µl) were placed on a cleaned, BSA-blocked coverslip in a custom cassette holder under humidification. Raw burst data was continuously acquired at a liner flow rate of 500 µm/sec with a sampling time of 500 µsec/data point. Unless otherwise indicated, single color BAS experiments with RuBisCO-A647 employed 50 µW of 642 nm laser excitation (at the sample). For single color BAS measurements using PepQ-TMR, 50 µW of 561 nm laser excitation (at the sample) was used. For MC-BAS experiments involving IbpA-OG or IbpBA488 and RuBisCO-A647, excitation laser power was experimentally calibrated in order to match the effective brightness of the dye pair in use: 150 µW 488 nm, IbpA-OG; 50 µW 488 nm, IbpB-A488; 50 µW 642 nm, RuBisCO-A647 when paired with IbpA-OG and 60 µW when paired with IbpB-A488. Burst detection, BAS and MC-BAS data analysis were carried out as previously described (40, 41).

## ACKNOWLEDGEMENTS

We would like to Dr. Chavela Carr for editorial contributions to the manuscript and members of the Rye lab for their valuable suggestions and discussions.

## AUTHOR CONTRIBUTIONS

Conceptualization: AR, JP, HSR

Methodology: JP, HSR

Investigation: AR

Formal Analysis: AR, JP

Validation: AR, JP

Supervision: HSR

Funding Acquisition: HSR

Writing - Original Draft: AR, HSR

Writing - Review and Editing: AR, JP, HSR

## DATA AVAILIBILITY

All data are available in the main text or the supplemental methods. Raw data sets available upon reasonable request from the corresponding author.

## SUPPLEMENTAL INFORMATION

**Supplemental Figure 1.**
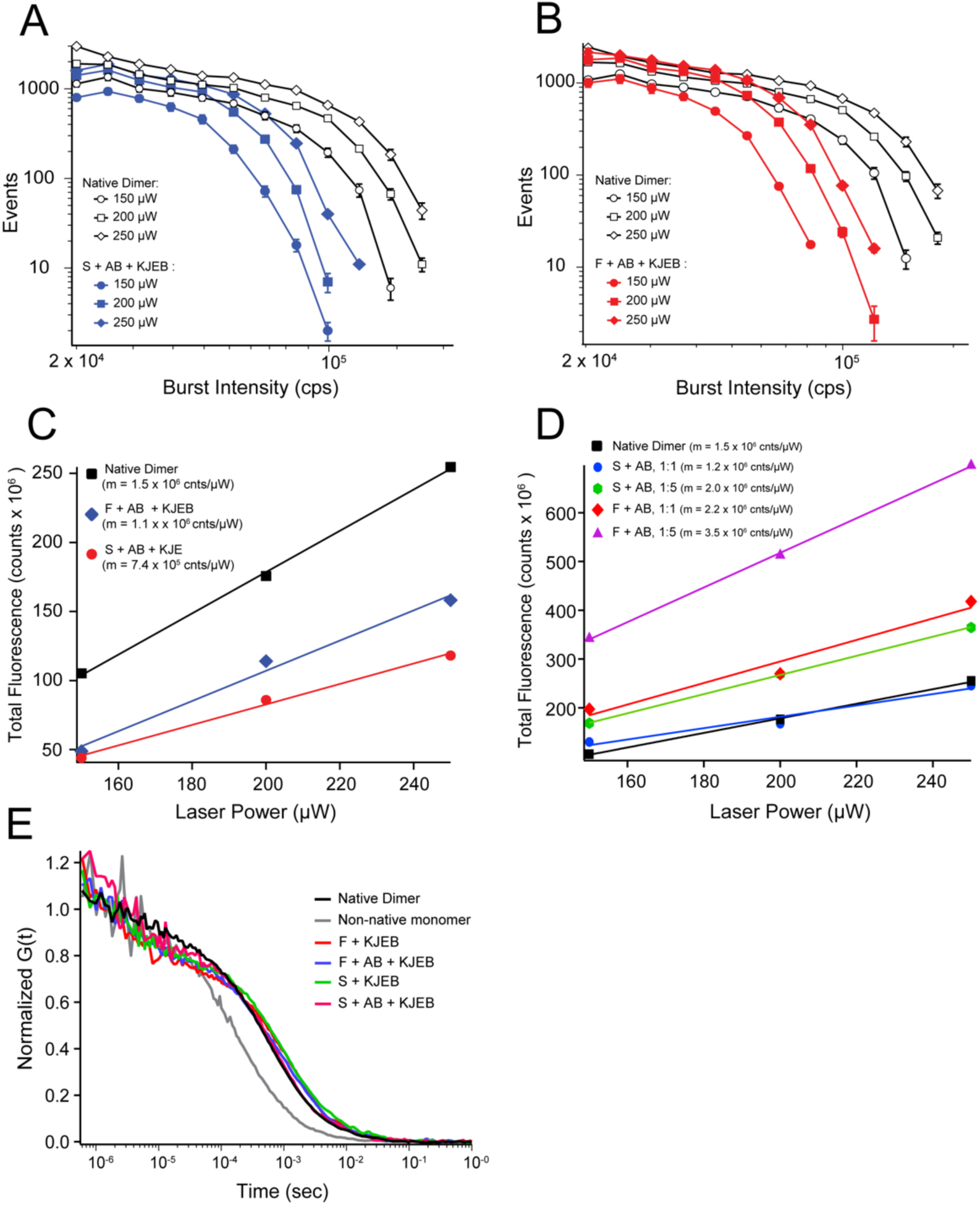
Calibration of intrinsic brightness for RuBisCO-A647. (A) Photon burst distribution of single, A647-labeled native RuBisCO dimers compared to a highly diluted endpoint of a KJEB-disassembly reaction of A647-labeled RuBisCO S-type aggregates containing IbpAB. Sample preparation and disassembly were accomplished as described in Figure 1 and 4. Each sample (< 50 pM labeled particles) flowed through the BAS microscope probe volume at a linear rate of 500 µm/sec. Samples were examined at three different excitation powers (642 nm laser) and each curve represents the average of three independent experiments, with the error bars displaying the standard deviation. The buffer background showed no detectable burst events above the minimal burst amplitude threshold employed (not shown). (B) Photon burst distribution of single, A647-labeled native RuBisCO dimers and a highly diluted disassembly endpoint of A647-labeled RuBisCO F-type aggregate particles containing IbpAB. Sample preparation and disassembly were accomplished as described in Figure 1 and 4 and data collection and analysis was accomplished as described for (A). (C) Dependence of total integrated fluorescence from single burst photon distribution curves (from A and B) on excitation laser power. (D) Dependence of total integrated fluorescence from single burst photon distribution curves (similar to A and B) for A647-labeled RuBisCO aggregate particles formed in the presence of different amounts of IbpAB: S + AB 1:1, S + AB 1:5, F + AB 1:1, and F + AB 1:5. (E) Normalized fluorescence correlation spectra (FCS) of a ∼ 100 pM sample of native RuBisCO dimer, a non-native RuBisCO monomer (41), and aggregate disassembly endpoints with or without IbpAB (A647 labeling in all cases).

**Supplemental Figure 2.**
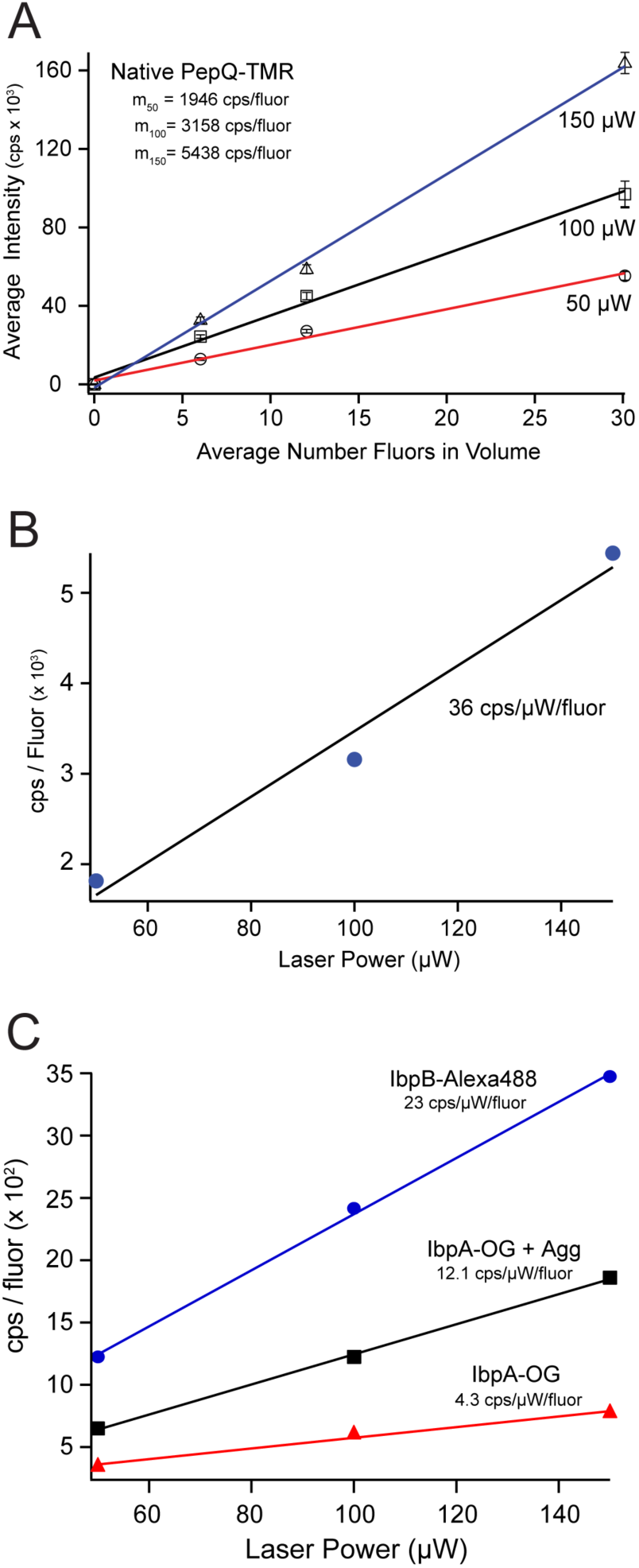
Calibration of intrinsic brightness for PepQ-TMR and labeled IbpAB variants. (A) Average fluorescence of known concentrations of native PepQ-TMR at different excitation laser powers (561 nm). Error bars display the standard deviation of three independent technical replicates. Fluorescence was collected using the same BAS microscope employed for all other experiments, with the same sample configuration, but in the absence of advective flow. (B) Dependence of the average fluorescence signal of PepQ-TMR on laser power. The slope of the lines gives the intrinsic brightness of the TMR dye on a native PepQ monomer (in counts per second / µW) for the microscope configuration used in all experiments. (C) Average fluorescence of known concentrations of IbpB-A488 and IbpA-OG, in the presence and absence of unlabeled RuBisCO aggregates (with and without S-type RuBisCO aggregates). In all cases, labeled IbpA or IbpB subunits are stoichiometrically paired with their unlabeled partner subunits. Fluorescence was collected using the same BAS microscope employed for all other experiments, with the same sample configuration, but in the absence of advective flow and following the same procedure used for panels (A) and (B). The slopes of each line give the intrinsic brightness of each probe in each context (in counts per second / µW).

**Supplemental Figure 3.**
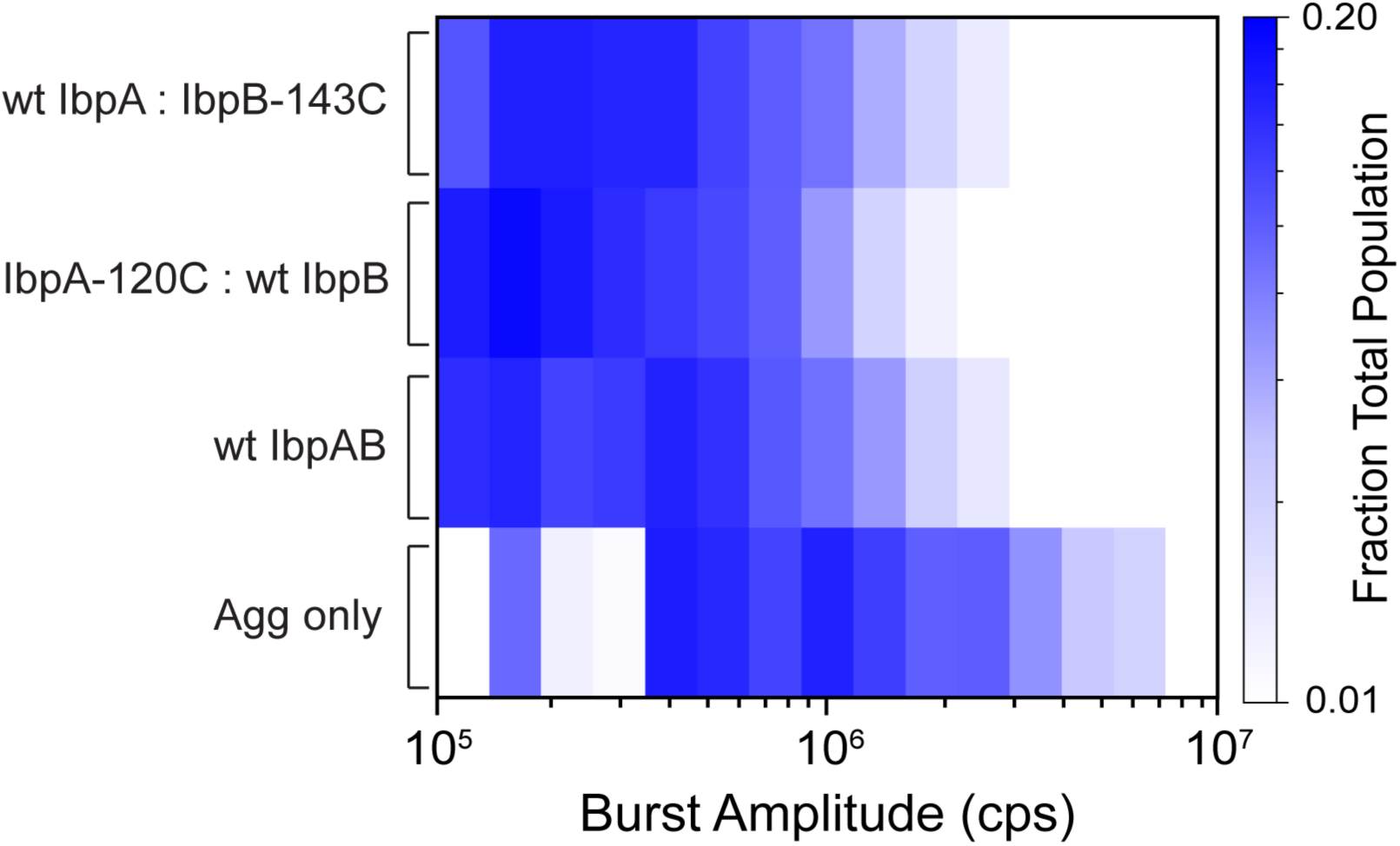
Inhibition of RuBisCO aggregation is not affected by IbpA-D120C and IbpB-143C mutations. The shift in the particle size distribution of S-type RuBisCO aggregates in the presence of wild-type and mutant IbpAB dimers was examined by BAS. In all cases, 200 nM Alexa647-labeled RuBisCO monomers were mixed with either buffer alone or heat-activated IbpAB (1:1 IbpAB:RuBisCO monomers), incubated for 2 min at 4 °C followed by incubation at 23 °C for 5 min, diluted 20-fold to halt particle growth and then examined by BAS. Particle size distributions for S-type aggregates alone (Agg only), wild-type IbpAB (wt IbpAB), IbpA-120C plus wild-type IbpB (IbpA-120C:IbpB) and wild-type IbpA plus IbpB-143C (wt IbpA:IbpB-143C) are shown. The BAS plot is a combination of n = 3, independent experimental replicates for each condition.

**Supplemental Figure 4.**
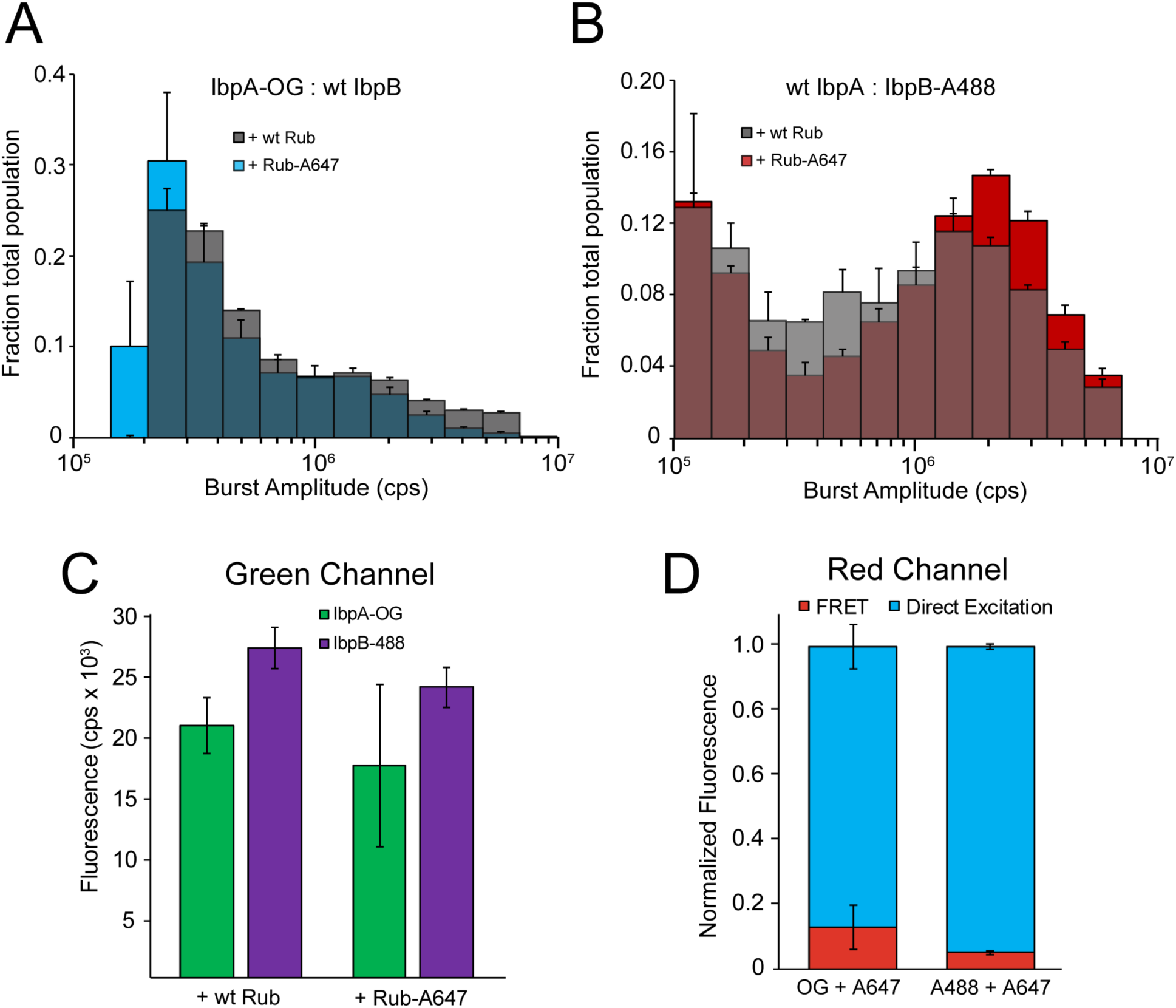
Co-aggregate particle distributions formed by fluorescently labeled IbpA and IbpB are minimally impacted by either RuBisCO labeling or FRET. (A) The size distribution of fluorescent particles containing IbpA-OG (A) or IbpB-A488 (B), formed in the presence of unlabeled (wt Rub, gray) or Alexa-647-labeled RuBisCO (Rub-A647, blue or red) was examined by BAS. IbpA-OG was mixed (1:1) with wild type IbpB and wild-type IbpA was mixed (1:1) with IbpB-A488. IbpAB mixtures were then heat activated, mixed (1:1) with 200 nM RuBisCO prepared for the S-type aggregation pathway and incubated for 2 min at 4 °C, followed by incubation at 23 °C for 5 min prior to BAS data collection. BAS distributions were calculated using bursts from the microscope blue detection channel only (488 nm excitation, 525 ± 18 nm emission). (C) The total integrated burst signal from the blue detection channel (6 min per record) is shown for aggregate particles containing either IbpA-OG (green) or IbpB-A488 (purple), formed with either unlableld or A647-labeled RuBisCO. (D) The total integrated burst signal from the microscope red detection channel (705 ± 36 nm emission) is shown for aggregate particles formed from A647-labeled RuBisCO, in the presence of IbpAB mixtures containing either IbpA-OG or IbpB-A488. For this measurement, either the 488 nm laser alone (direct excitation of either the OG or A488 probe) or 642 nm laser alone (direct excitation of the A647 probe) was employed. The red bar illustrates the average signal in the red detection channel attributable to energy transfer (FRET) between the OG or A488 (as donors) and the A647 probe (as acceptor) during an MC-BAS experiment. The blue bar illustrates the average signal in the red detection channel attributable to direct excitation. All data shown is derived from a combination of n = 3, independent experimental replicates and error bars show ± s.d.

**Supplemental Figure 5.**
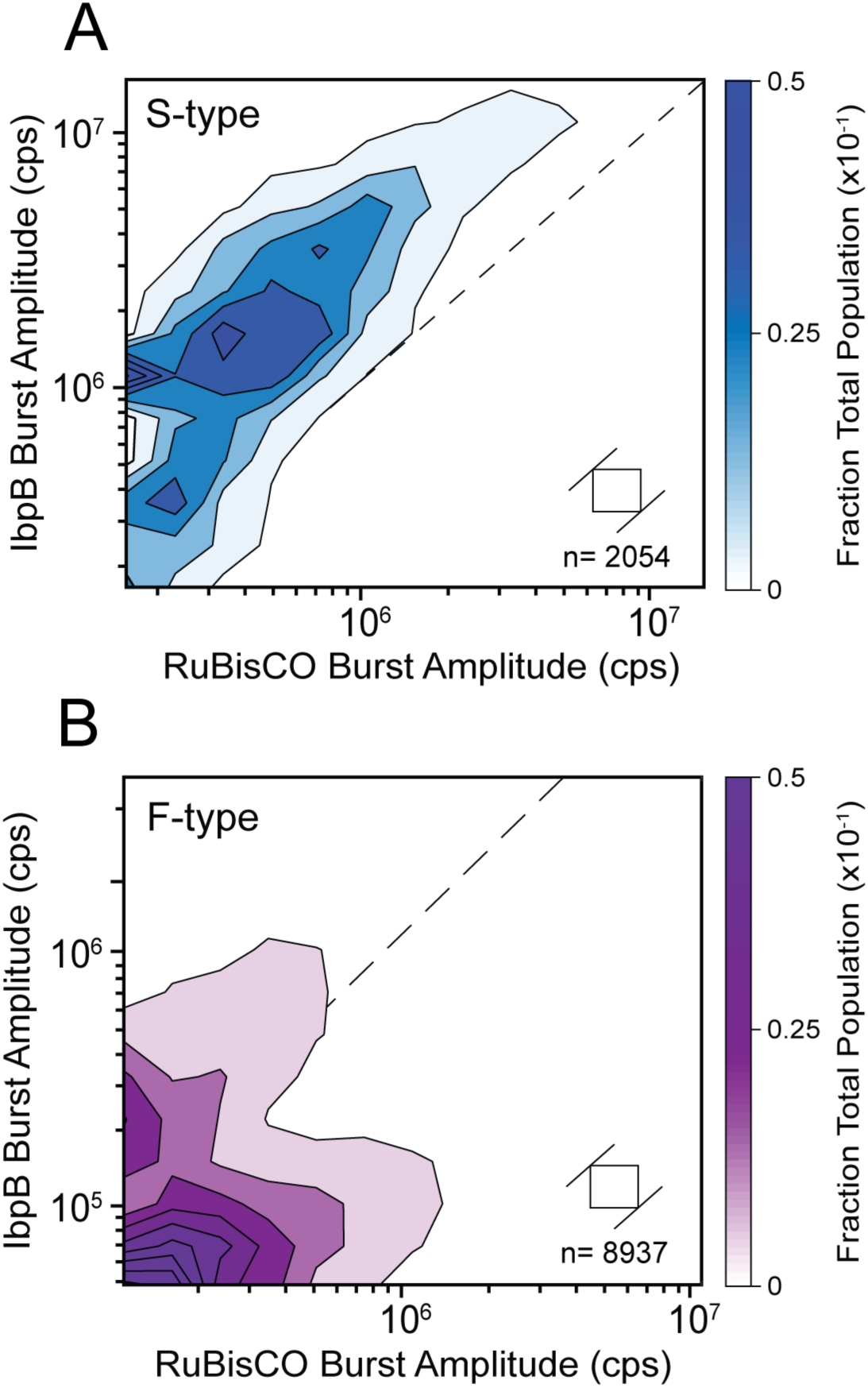
The binding distribution of IbpAB on RuBisCO aggregates observed with labeled IbpB. The MC-BAS distributions for (A) S-type and (B) F-type RuBisCO aggregates bound to IbpAB using IbpB-A488. In each case, the final mixing ratio of RuBisCO monomers to IbpAB dimers was 1:1. The RuBisCO burst intensity is plotted on the x-axis and IbpAB burst intensity is plotted on the y-axis. The dashed diagonal line shows the experimentally determined 1:1 brightness equivalence for the RuBisCO- and IbpB-coupled A488 dye. The spread of the distributions along the positive diagonals of the plot measures the population size distribution at a given IbpAB:RuBisCO stoichiometry, while the extent of spread along the negative diagonals is proportional to the range of binding stoichiometries. Each MC-BAS plot is a combination of n = 6, independent experimental replicates. The square in each plot shows the 2D bin size prior to contour plot extrapolation and n indicates the total number of coincident burst events in data set.

**Supplemental Figure 6.**
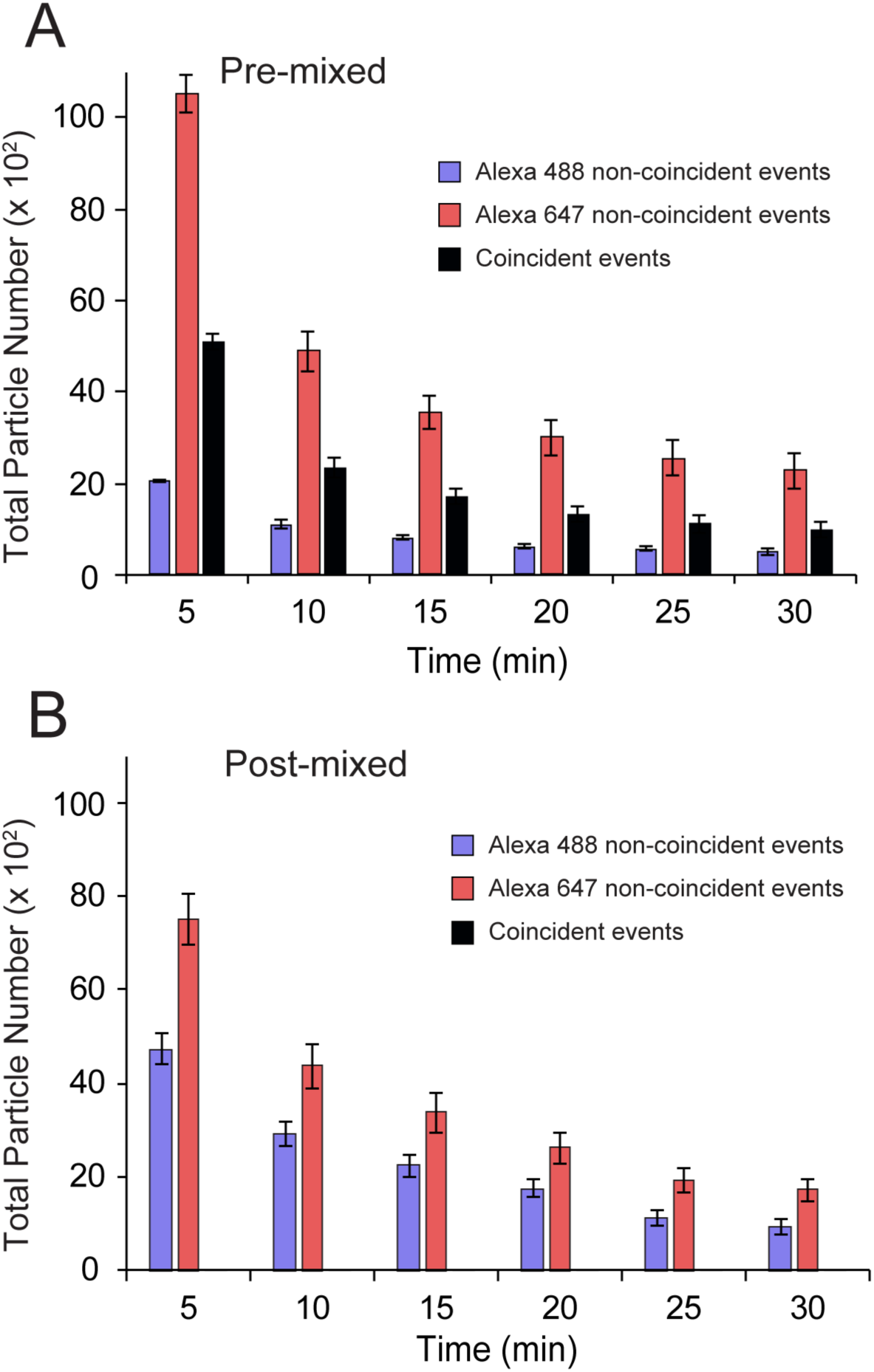
IbpAB-bound RuBisCO aggregate particles do not reform during disaggregation. (A) Disassembly of co-labeled aggregate particles observed by MC-BAS as a function of time, compared with the number of particles that are only detectable in one or the other channel. S-type aggregates were formed in the presence of IbpAB at a mixing ratio of 1:1, using a 1:1 mixture of two, differently labeled RuBisCO monomers, one carrying a single Alexa488 dye and the other a single Alexa647 dye (pre-mixed). Following formation of co-labeled aggregates (10 nM final total monomer), aggregates were supplemented with 250 nM DnaK, 500 nM DnaJ, 500 nM GrpE, 50 nM ClpB, 2 mM ATP, and a creatine kinase-based ATP regeneration system. (B) Disassembly of aggregate particles by MC-BAS in which each labeled monomer was separately aggregated with IbpAB at a mixing stoichiometry of 1:1, supplemented with the KJEB system and then co-mixed after 10 sec (post-mixed). Aggregation conditions, final total monomer and KJEB component concentrations are identical to the pre-mixed experiment in panel (A). For post-mixed samples, no particles displaying significant co-incident signals in both channels could be detected over the course of the disassembly experiment. Error bars display the standard deviation of the total number of particles detected at each time point from n = 3, independent experimental replicates.

**Supplemental Figure 7.**
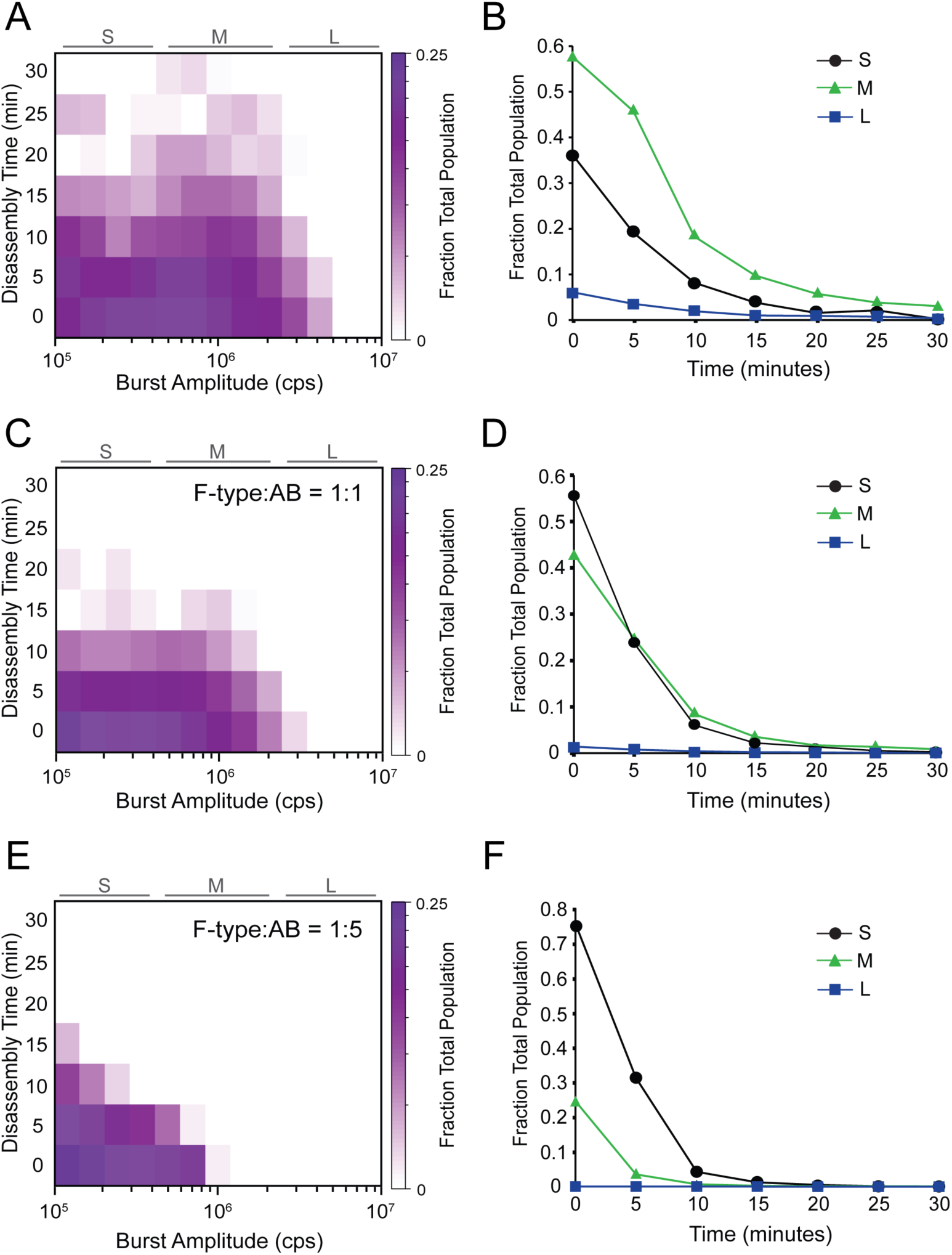
IbpAB accelerates the disassembly of F-type RuBisCO aggregates by the KJEB bi-chaperone disaggregase. F-type aggregates were formed in either the absence (A) or presence of IbpAB at two different RuBisCO monomer to IbpAB ratios: 1:1 (C) and 1:5 (E). Disaggregation was triggered by the addition of the KJEB bi-chaperone system (1 µM DnaK, 2 µM DnaJ, 2 µM GrpE and 200 nM ClpB), 2 mM ATP, and a creatine kinase-based ATP regeneration system. Samples were then loaded onto a BAS microscope and burst data was continuously acquired for 30 min. The full experimental photon history was segmented into 5 min blocks and analysis was performed on each block. The heat maps represent a combination of three, independent experimental replicates for each aggregation condition. A zero-time measurement on each sample was collected prior to the addition of ATP. To highlight how disaggregation rates are impacted by aggregate size, the BAS heat maps were also coarsely binned into small (*S*), medium (*M*), and large (*L*) particle ranges (B, D, and F). In each case, all detected objects within a given size range were summed and plotted as a function of time following the initiation of disaggregation by KJEB.

**Supplemental Figure 8.**
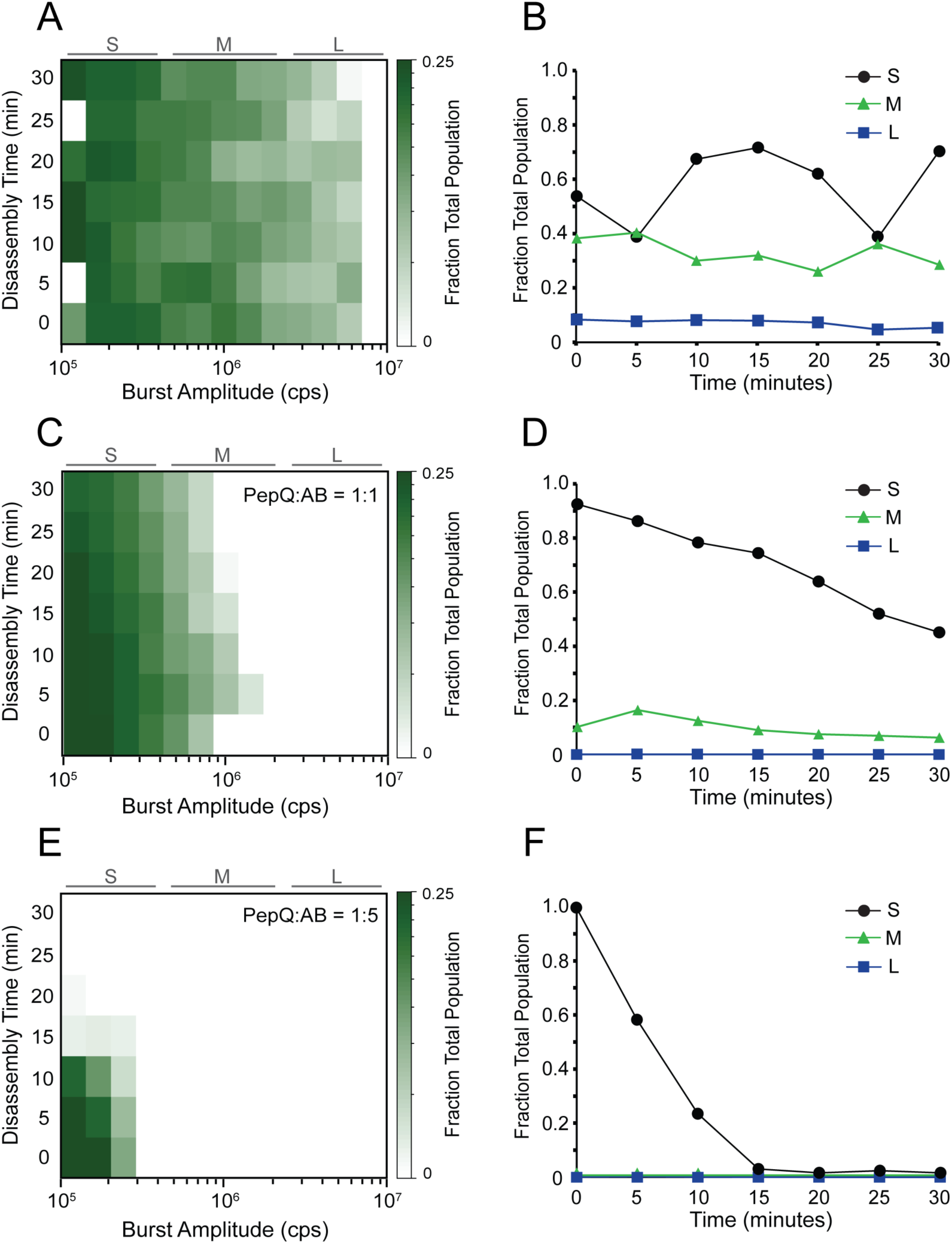
IbpAB dramatically accelerates the disassembly of PepQ aggregates by the KJEB bi-chaperone disaggregase. PepQ aggregates were formed in either the absence (A) or presence of IbpAB at two different PepQ monomer to IbpAB ratios: 1:1 (C) and 1:5 (E). Disaggregation was triggered by the addition of the KJEB bi-chaperone system (1 µM DnaK, 2 µM DnaJ, 2 µM GrpE and 200 nM ClpB), 2 mM ATP, and a creatine kinase-based ATP regeneration system. Samples were then loaded into a BAS microscope and burst data was continuously acquired for 30 min. The full experimental photon history was segmented into 5 min blocks and analysis was performed on each block. The heat maps represent a combination of n = 3, independent experimental replicates for each aggregation condition. A zero-time measurement on each sample was collected prior to the addition of ATP. To highlight how disaggregation rates depended on aggregate size, the BAS heat maps were also coarsely binned into small (*S*), medium (*M*), and large (*L*) particle ranges (B, D, and F). In each case, all detected objects within a given size range were summed and plotted as a function of time following the initiation of disaggregation by KJEB.

**Supplemental Figure 9.**
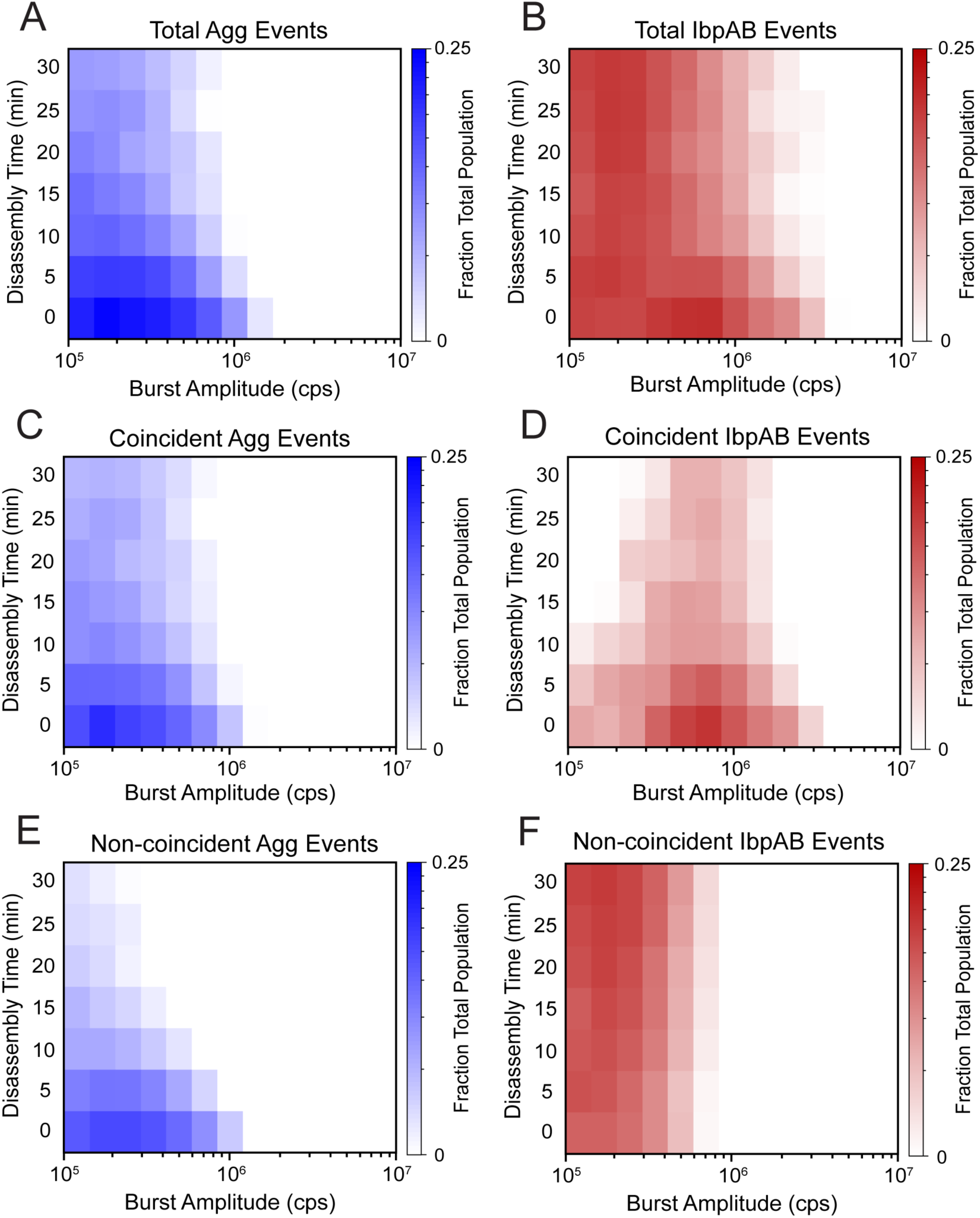
Disassembly of IbpAB-bound RuBisCO aggregate particles slows when KJEB levels are reduced. The data for each channel of the MC-BAS experiment shown in Figure 6 were separately examined by standard BAS. S-type RuBisCO-A647 aggregates, formed in the presence of IbpAB (IbpA-OG) at a mixing ratio of 1:1 RuBisCO:IbpAB, were subjected to disassembly at a reduced KJEB concentration (250 nM DnaK, 500 nM DnaJ, 500 nM GrpE and 50 nM ClpB). The photon histories from the (A) RuBisCO and (B) IbpAB channels were segmented into five-minute bins and each temporal bin was then examined by BAS to give the population-resolved kinetics of total RuBisCO and IbpAB particle disassembly. The population-resolved kinetics of the coincident events only (C and D), as well as the non-coincident events only (E and F), were also examined.

**Supplemental Figure 10.**
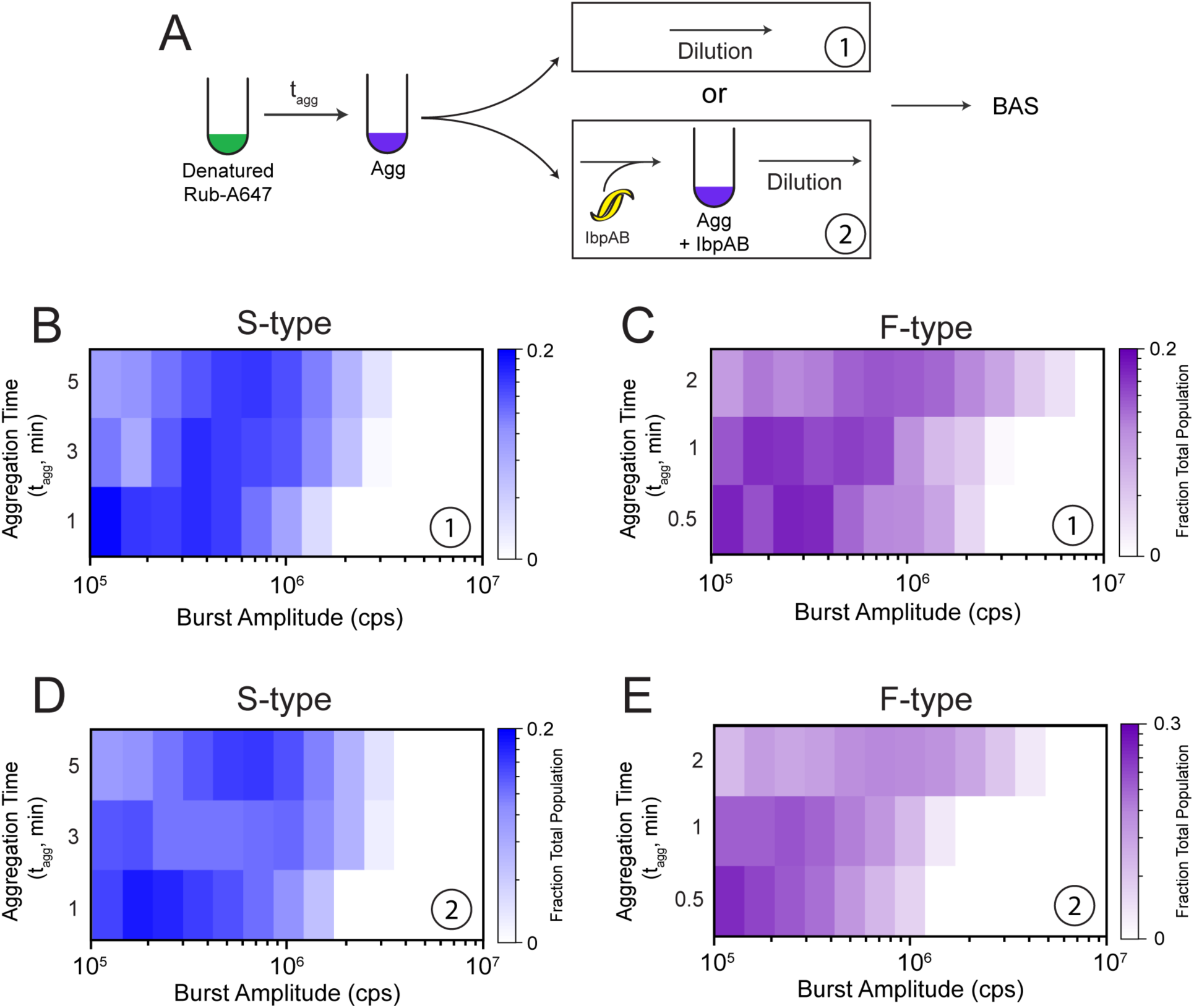
Delayed addition of IbpAB halts, but does not reverse, aggregate particle growth. (A) Experimental protocol for examining the consequences of delayed IbpAB addition. At different times (t_agg_) following the initiation of aggregation, samples were either directly diluted to halt aggregation (1) or mixed with IbpAB at 1:1 and incubated for additional time prior to dilution and measurement (2). S-type aggregate samples were incubated for a total of 5 min prior to BAS measurement and fibril-like samples were incubated for a total of 2 min. The population-resolved kinetics of particle growth for S-type (B) and F-type (C) aggregates in the absence of IbpAB are shown. Delayed addition of IbpAB to either S-type (D) or F-type (E) aggregates halts particle growth at approximately the same particle distribution as dilution alone, with little or no apparent additional change in the particle size distribution over time. Each BAS plot is a combination of n = 3, independent experimental replicates.

